# PIWIL3-piRNA Pathway Is Essential for Rabbit Oogenesis and Embryogenesis via Broad Regulation of the Transcriptome and Proteome

**DOI:** 10.1101/2025.10.23.684072

**Authors:** Yuanyuan Gong, Ling Li, Yuqiang Qian, Ting Lu, Zhaoran Zhang, Lichun Jiang, Guohui Liu, Meihua Cui, Shangang Li, Zhanjun Li, Shuo Shi, Haifan Lin

## Abstract

Female infertility represents a significant challenge in reproductive medicine. Although it is known to be primarily due to oogenic and early embryonic failures, the molecular mechanisms underlying these failures remain elusive. The Piwi-piRNA pathway is crucial for gametogenesis in diverse organisms. Yet, its role in mammalian female fertility is unclear. This is partly because most studies are done in mice, which contain only PIWIL1, PIWIL2, and PIWIL4 that are essential for spermatogenesis but not for oogenesis. PIWIL3 emerges in higher mammals and is highly expressed in human oocytes. Interestingly, female *PIWIL3*-knockout golden hamsters exhibit only reduced fertility without detectable defects in oogenesis. This has left the function of PIWIL3 in higher mammals, including humans, unexplored. Here, we discovered that PIWIL3 in the rabbit (*Oryctolagus cuniculus*) shares high homology with human PIWIL3. It is the predominant PIWI protein in oocytes in both species. Using CRISPR–cas9-mediated knockout, we demonstrated that rabbit PIWIL3 is essential for female fertility. Its loss leads to severe defects in oogenesis. Moreover, embryos lacking maternal PIWIL3 arrest developmentally at the 8-cell stage. Mechanistically, rabbit PIWIL3 binds ∼18-nucleotide piRNAs, mirroring the behavior of human PIWIL3, and is critical for piRNA biogenesis. Moreover, it regulates transcriptomic and proteomic landscapes and silences a broad array of transposons during late oogenesis yet activates another set of transposons during early embryogenesis. These findings establish PIWIL3 as a pivotal dual regulator of gene expression and transposon activity, which is essential for oogenesis and early embryogenesis in non-rodent mammals, potentially including humans.

## INTRODUCTION

In humans and other mammals, defects in oocyte number and quality, as well as in early embryogenesis are primary causes of female infertility—a persistent clinical and scientific challenge^1, 2^. The generation of developmentally competent oocytes requires intricately orchestrated molecular events spanning transcriptional, post-transcriptional, and translational regulatory layers^3–5^. These complex programs precisely coordinate oocyte growth, meiotic maturation, and the subsequent embryonic development following fertilization^6–14^. Disruption of these regulatory networks can diminish ovarian reserve, impair oocyte quality, and lead to embryonic developmental arrest^9–14^. Despite significant progress, the master regulators responsible for coordinating multi-level gene expression dynamics during the oocyte-to-embryo transition (OET) remain incompletely understood.

The PIWI-piRNA pathway is a crucial regulator of germline biology, initially recognized for silencing transposable elements (TEs) to safeguard genomic integrity^15–20^. PIWI proteins bind Piwi-interacting RNAs (piRNAs), forming piRNA-induced silencing complexes (piRISCs) that mediate transcriptional and post-transcriptional gene regulation^21–32^. Beyond TE repression, this pathway also regulates protein-coding and non-coding RNAs during embryogenesis, tissue regeneration, and even oncogenesis^33–41^. However, functional diversification among *PIWI* paralogues has created species-specific regulatory landscapes, complicating generalization across mammals. For instance, mice—the predominant mammalian model—possess three *PIWI* genes (*PIWIL1*, *PIWIL2*, and *PIWIL4*) that are essential for spermatogenesis but have no detectable role in oogenesis or female fertility, either individually or collectively ^42–44^. However, an additional *PIWI* gene, *PIWIL3*, emerges in higher mammals. Recent studies in golden hamsters (*Mesocricetus auratus*), another rodent species, have revealed that *PIWIL1*-deficiency causes complete female sterility whereas *PIWIL3*-deficiency reduces female fertility by affecting early embryogenesis of progeny^30, 31, 45^, indicating an important role of PIWIL1 and largely redundant role of *PIWIL3* in female reproduction. Notably, neither *PIWIL1* or *PIWIL3* has detectable function in oogenesis; especially, PIWIL3 has little effect even on gene expression and transposon activity^30,45^.

Species-specific differences are further exemplified by piRNA populations within mammalian oocytes. Hamster oocytes harbor three distinct piRNA classes: 22–24 and 28-30 nucleotide (nt) piRNAs bound by PIWIL1 and an 18-20 nt piRNA subset bound by PIWIL3^30–32, 45^. By contrast, human oocytes predominantly express 18-21 nt piRNAs, which are primarily bound by PIWIL3^28, 31, 46^. Correspondingly, *PIWIL3* mRNA and protein are highly expressed as the predominant *PIWI* member in human oocytes^27, 31, 47^, whereas *PIWIL1* is the principal *PIWI* in hamster oocytes^29, 31^. Overall, this divergence presents an evolutionary paradox: although PIWIL3 is conserved across mammals, it exhibits only minor functional significance in female rodent reproduction, limiting the direct translation of rodent findings to human biology.

Addressing this translational gap requires identifying a model organism whose PIWI-piRNA dynamics more accurately reflect those in humans. We hypothesized that the phylogenetic proximity of certain species could provide a viable solution. The New Zealand White Rabbit (*Oryctolagus cuniculus*) emerged as a promising candidate based on two key factors: (1) Phylogenetic analysis reveals a closer evolutionary relationship between rabbit *PIWI* subfamily members and humans as compared to rodents. Notably, rabbit *PIWIL3* shows a closer evolutionary distance to humans even than that of cattle and pigs, second only to rhesus monkeys (*Macaca mulatta*) (Supplementary information, Fig. S1a). (2) Tissue-specific expression profiling reveals a pronounced ovarian enrichment of PIWIL3 in rabbits, mirroring the expression pattern observed for human PIWIL3 ^27, 31, 45, 47^, whereas PIWIL1 and PIWIL2 remain testis-specific (Fig. 1a). These features position the rabbit as a compelling model to investigate the roles of *PIWIL3* in female gametogenesis and early development, bridging the gap between rodent models and human reproductive biology.

**Fig. 1.**
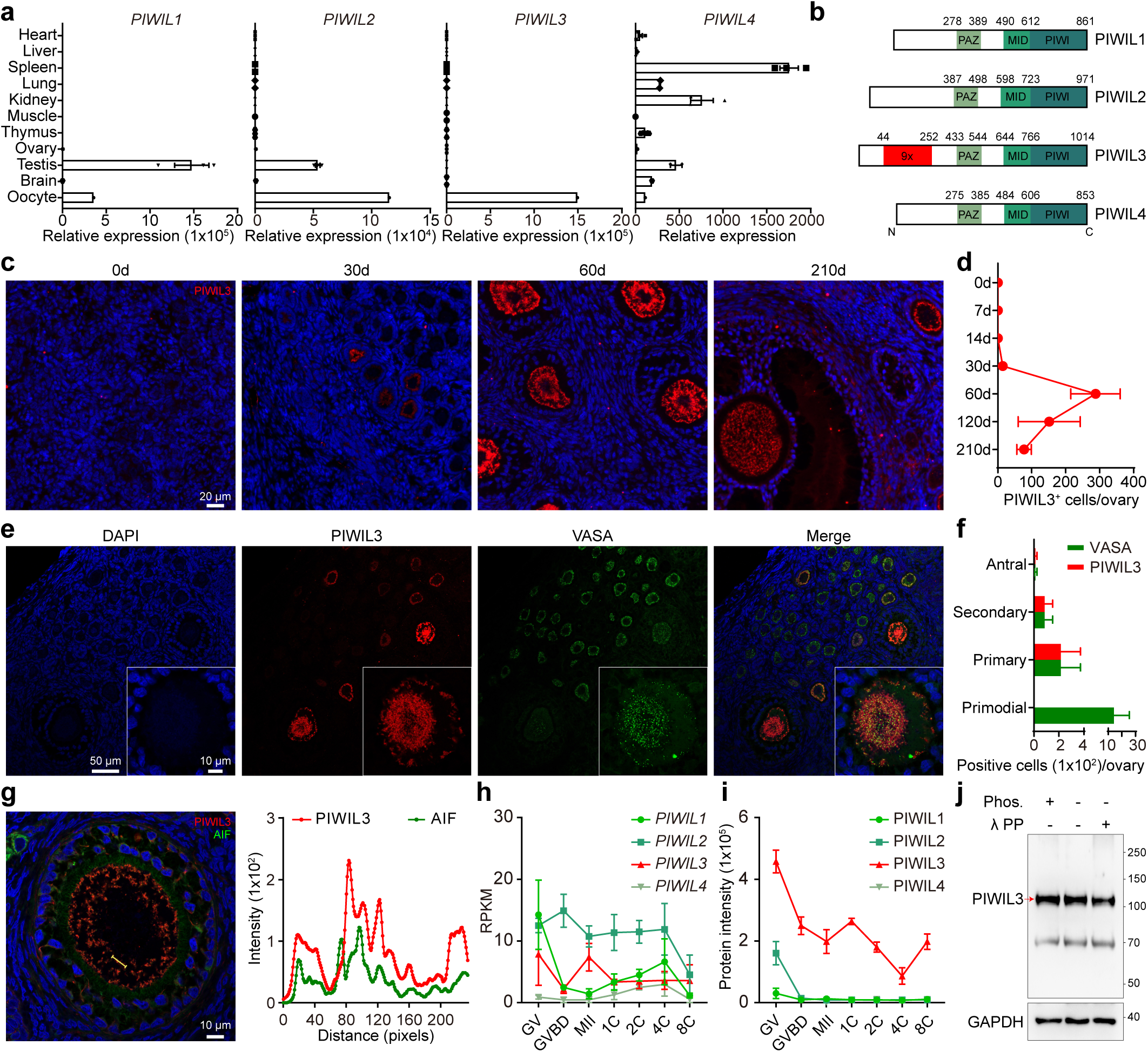
PIWIL3 is highly expressed in growing oocytes and early embryos of rabbits. **a** qRT-PCR analysis of mRNA expression levels of four *PIWI* genes (*PIWIL1*, *PIWIL2*, *PIWIL3*, and *PIWIL4*) in various rabbit tissues. *β-Actin* was used as an internal control to normalize the expression levels of *PIWI* genes across samples. Data are presented as the mean ± SD from three independent experiments. **b** Protein amino acid sequence alignment of four *PIWI* genes in rabbits. The red regions represent nine repeated nucleotide sequences that encode the amino acid motif GAAWERG. **c** PIWIL3 immunofluorescence staining of ovarian sections from rabbits of different ages (0 days, 30 days, 60 days, and 210 days of age). Scale bars = 20 µm. **d** Line graph showing the quantitative analysis of PIWIL3-positive-staining cells in ovarian sections from rabbits of different ages. Data are presented as the mean ± SD from three independent ovarian sections. **e** Co-immunofluorescence staining of PIWIL3 (red) and VASA (green) in 210-day-old rabbit ovarian sections. Scale bars = 50 µm (left) and 10µm (right). **f** Bar graph showing the quantitative analysis of PIWIL3 and VASA co-positive staining cells in different follicle types in 210 days rabbit ovarian sections. **g** Representative co-immunostaining of PIWIL3 (red) and mitochondrial marker AIF (green) in rabbit oocytes (left). Scale bars = 10 µm, intensity profiles of PIWIL3 and AIF in representative image along the yellow line (right). **h** mRNA expression levels of *PIWI* genes in rabbit oocytes (GV, GVBD, MII) and embryos (1C, 2C, 4C, 8C), normalized as reads per kilobase of transcript per million mapped reads (RPKM). **i** Protein expression levels of *PIWI* genes in rabbit oocytes (GV, GVBD, MII) and embryos (1C, 2C, 4C, 8C), normalized as protein intensity. **j** Western blotting was performed from rabbit MII oocytes with (+) or without (−) treatment of phosphatase inhibitor (Phos.) and λ protein phosphatase (λ PP). The red arrow indicates the PIWIL3 protein band, with an estimated molecular weight of 110 kD(A) GAPDH was used as the loading control.

Using CRISPR/Cas9-mediated knockout and multi-omics analyses, we demonstrate the critical and pleiotropic roles of PIWIL3 during rabbit oogenesis and early embryogenesis. Our findings reveal that PIWIL3 orchestrates transcriptomic and proteomic reprogramming, is indispensable for piRNA biogenesis, and dynamically regulates transposable element expression throughout the OET. These insights significantly expand our understanding of PIWIL3 function beyond established rodent models and provide a comprehensive framework to investigate PIWI-piRNA pathway dysregulation in human female infertility.

## RESULTS

### *PIWIL3* is highly expressed in growing oocytes and early embryos of rabbits

To begin our investigation of PIWIL3 gene in rabbits, we first analyzed its expression profile across various rabbit tissues using quantitative Reverse Transcription Polymerase Chain Reaction (qRT-PCR). Our results indicated that PIWIL3 was specifically expressed in the oocyte, whereas the other three *PIWI* genes exhibited broader expression patterns, detected not only in oocytes but also in the testis and several other tissues (Fig. 1a). This finding underscores a unique role for PIWIL3 during rabbit oogenesis.

We next cloned ovarian cDNAs encoding *PIWIL3* and characterized their sequences, identifying four distinct isoforms expressed in rabbit ovaries (Supplementary information, Fig. S1b). Notably, these isoforms exhibited variations in their N-terminal regions compared to the sequences predicted by the Ensembl and UniProt databases. Isoform 1, the longest variant, had a coding sequence (CDS) comprising 20 exons and encoding a 1,014-amino acid protein that featured nine repeats of the nucleotide sequence encoding the GAAWERG motif. Isoform 2 consisted of a CDS of 19 exons encoding an 810-amino acid protein. Similarly, Isoform 3 also contained 19 exons but lacked a substantial portion of exon 3 found in Isoform 1, resulting in a shorter protein of 762 amino acids. Isoform 4 was the shortest, with a CDS of 13 exons encoding a protein of 537 amino acids.

To further characterize PIWIL3 protein expression and localization during rabbit ovarian development, we generated mouse monoclonal antibodies targeting the PIWIL3 protein (see Methods). Immunostaining demonstrated initial detection of PIWIL3 in ovarian tissues from 30-day-old rabbits, with PIWIL3-positive cells peaking in number at 60 days (Fig. 1c, d). Additionally, co-localization analyses with the germ cell marker VASA revealed specific expression of PIWIL3 in the cytoplasm of growing oocytes within primary and secondary follicles (Fig. 1e, f). Moreover, co-localization studies with the mitochondrial marker Apoptosis-inducing factor (AIF) showed abundant localization of PIWIL3 protein to mitochondria (Fig. 1g).

We then analyzed the mRNA and protein expression patterns of *PIWIL3* and other *PIWI* genes throughout oocyte maturation stages (germinal vesicle (GV) to metaphase II (MII)) and during early embryonic development (from the 1-cell (1C) to 8-cell (8C) stages) (Fig. 1h, i). While *PIWIL3* mRNA expression levels did not significantly exceed those of other PIWI genes in oocytes and early embryos, mass spectrometry analysis demonstrated substantially higher expression of PIWIL3 protein relative to other PIWI proteins in GV-stage oocytes. Importantly, upon entry into germinal vesicle breakdown (GVBD), PIWIL3 became the sole PIWI protein detected in oocytes (Fig. 1i). This exclusive presence of PIWIL3 persisted through the MII stage and early embryogenesis, indicating a robust translational repression of all *PIWI* family mRNAs except *PIWIL3* during these developmental periods.

Considering previous reports highlighting significant phosphorylation of PIWIL3 protein in golden hamster MII oocytes^29^, we treated rabbit MII oocyte lysates with calf intestinal alkaline phosphatase followed by SDS electrophoresis and western blotting to detect any potential down-shift of PIWIL3 due to dephosphorylation. However, the position of the PIWIL3 protein band remained unchanged at approximately 110 kDa (Fig. 1j). These data suggest that Isoform 1 is the predominant variant expressed in rabbit oocytes and that rabbit PIWIL3 protein is not detectably phosphorylated at the MII stage (Fig. 1j).

### *PIWIL3*-deficient female rabbits display oogenic and maternal-effect embryogenic defects, leading to complete infertility

To investigate the role of PIWIL3 in rabbit female reproduction, we generated two *PIWIL3*-knockout rabbit mutants from two different rabbit strains using CRISPR-Cas9-mediated pronuclear injection at day 5, targeting exon 5 of the *PIWIL3* gene with dual guide RNAs (Fig. 2a; Supplementary information, Fig. S2a). The *PIWIL3*^1/1^ mutant carried a 5-nt deletion and the *PIWIL3*^2/2^ mutant harbored a 178-nt deletion (Supplementary information, Fig. S2a, b). Both mutations introduced frameshifts that led to premature stop codons in *PIWIL3* mRNA (Supplementary information, Fig. S2c). In homozygous *PIWIL3* mutants, PIWIL3 protein was undetectable in MII oocytes of both mutants, as confirmed by western blotting (Supplementary information, Fig. S2d) and immunostaining (Supplementary information, Fig. S2e). Notably, both homozygous *PIWIL3* mutant females were completely sterile (Fig. 2b, c). Despite successful copulation, mating these females with wild-type males resulted in neither pregnancy (Fig. 2b) nor litters (Fig. 2c), underscoring the strict maternal effect of *PIWIL3* on embryogenesis.

**Fig. 2.**
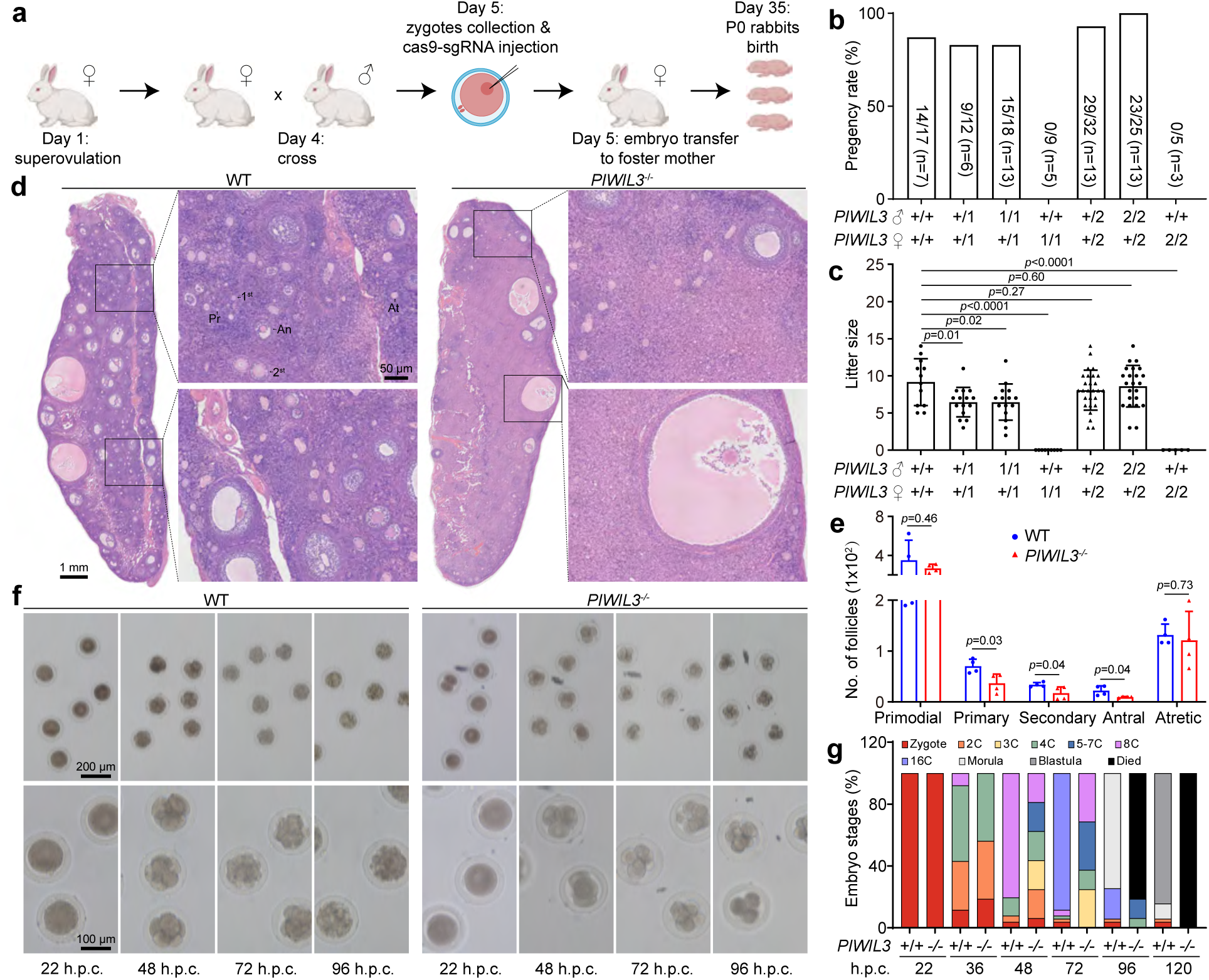
*PIWIL3* deficiency causes oogenesis defects and infertility of female rabbits. **a** Strategy for the generation of *PIWIL3*-deficient rabbits by pronuclear CRISPR-Cas9 injection. **b** The fecundity of *PIWIL3*-deficient female rabbits was assessed by natural mating; n represents the number of female rabbits for each genotype. The numbers on the bars indicate the pregnancy rate per coitus. **c** Litter sizes from mated wild-type (WT) and *PIWIL3* mutant (1 and 2 refer to two *PIWIL3* mutant strains) females. Error bars represent ± s.e.m., and statistical significance (*p* value) was assessed using unpaired two-tailed Student’s t-test. **d** Hematoxylin and eosin (H&E) staining of WT and *PIWIL3*-deficient (*PIWIL3^-/-^,* include *PIWIL3*^1^*^/^*^1^ and *PIWIL3*^2^*^/^*^2^) in 210-day-old rabbit ovarian sections. Primordial follicles (Pr), primary follicles (1°), secondary follicles (2°), antral follicles (An), and atretic follicles (At) represent stages of normal follicular development. Ovaries from three WT and *PIWIL3*-deficient female rabbits were analyzed, with representative images shown. Scale bars = 1 mm (left panel) and 50 µm (right panel). **e** Bar graph showing the quantification of different follicle types in ovarian sections from WT and *PIWIL3*-deficient rabbits. Error bars represent ± s.e.m., and statistical significance (*p*-value) was assessed using unpaired two-tailed Student’s t-test. **f** Representative images illustrating the delayed development of maternal *PIWIL3*-deficient embryos cultured *in vitro* at the indicated (22, 48, 72, and 96) hours post coitum (h.p.c.). Scale bars = 200 µm (upper panel) and 100 µm (lower panel). **g** Bar graph depicting the embryogenesis ratio of WT (n = 51) and maternal *PIWIL3*-deficient (n = 16) embryos in panel **f**.

Additionally, when *PIWIL3*^+/1^ females were crossed with either heterozygous or homozygous *PIWIL3*-knockout males, litter sizes were reduced, whereas *PIWIL3*^+/2^ females crossed with heterozygous or homozygous *PIWIL3*-knockout males showed no significant change in litter size (Fig. 2c). This differential dose sensitivity of PIWIL3 may reflect genetic background differences between the two mutant strains.

To identify specific defects in *PIWIL3* mutant females, we conducted a detailed analysis of oogenesis and early embryogenesis. Histological examination of ovarian sections from *PIWIL3*^1/1^ and *PIWIL3*^2/2^ rabbits revealed dramatic reductions in primary, secondary, and antral follicles containing growing oocytes, despite ovarian size and weight comparable to wild-type controls (Fig. 2d, e; Supplementary information, Fig. S2f, g). Intriguingly, at Day 0—prior to the onset of PIWIL3 protein expression— oogonia numbers were unaffected by PIWIL3 deficiency (Supplementary information, Fig. S2h, i), indicating that PIWIL3 plays a stage-specific role in oogenesis and oocyte maturation.

Because *PIWIL3* knockout ovaries still produced some mature oocytes, the complete infertility of the knockout females pointed to potential maternal effects on the subsequent embryogenesis of these oocytes after fertilization. To explore this possibility, we fertilized wild-type sperm with oocytes derived from either wild-type or *PIWIL3*-deficient females and monitored embryonic development. Maternally *PIWIL3*-deficient embryos exhibited developmental delays as early as 36 hours post-coitus (h.p.c.), with an increased proportion of 1-cell embryos and a complete absence of 8-cell embryos (Fig. 2f, g). By 48 h.p.c., 80.4% of wild-type embryos progressed to the 8-cell stage. However, only less than 20% of *PIWIL3*-deficient embryos reached this milestone, reflecting severe developmental delay. Furthermore, maternally *PIWIL3*-deficient embryos frequently arrested at intermediate stages (3-7 cells), in contrast to the small number of delayed wildtype embryos, which mostly at 4-cell stage, with a few at 2-cell or zygote stage. This indicates that maternal *PIWIL3*-deficiency causes asynchronous divisions. Developmental impairment became more pronounced at 72 h.p.c., when wild-type embryos advanced to the 16-cell stage, maternally *PIWIL3*-deficient embryos were still at 3-8-cell stages. By 96 h.p.c., when most wild-type embryos had formed morulae, maternally *PIWIL3*-deficient embryos were mostly dead, with only a few surviving but arrested at 5-7-cell stages. By 120 h.p.c., when wild-type counterparts successfully reached the blastula stage, all maternally *PIWIL3*-deficient embryos had perished (Fig. 2f, g). These findings demonstrate that *PIWIL3*-deficiency not only reduces ovarian oocyte reserve but also causes severe defects of the resulting embryos, leading to early embryogenic lethality.

In contrast, *PIWIL3*-knockout males had no fertility impairments. Their testicular morphology and histology were indistinguishable from wild-type controls (Supplementary information, Fig. S2j, k). These observations indicate that PIWIL3 is specifically required for female fertility in rabbits.

### PIWIL3 is required for transcriptomic reprogramming during the oocyte maturation and zygotic genome activation

The essential roles of Rabbit *PIWIL3* for oogenesis and early embryogenesis suggests that *PIWIL3* may play a prominent role in regulating the oocyte and early embryonic transcriptome in rabbits and potentially other higher mammals.

To investigate this role, we examined whether PIWIL3 regulates the dynamic transcriptomic changes occurring during OET and how this regulation influences oocyte maturation and embryonic development. Using Smart-seq3, we profiled transcriptomes from single oocytes and early embryos spanning the GV to the 16C stages. Principal component analysis (PCA) revealed distinct transcriptomic reprogramming events at successive stages. In controls (*PIWIL3^+/+^*or *PIWIL3^+/-^*), a pronounced shift occurred at the GV-to-GVBD transition, showing strong divergence between GV and GVBD transcriptomes (Fig. 3a). A subsequent, smaller shift was observed between GVBD and MII oocytes. Post-fertilization, transcriptomes remained relatively stable from MII through the 4C stage but underwent two other major phases of changes at the 4C-to-8C and 8C-to-16C transitions, indicating zygotic genome activation (ZGA) and a decline of maternal mRNA dominance after the 4C stage.

**Fig. 3.**
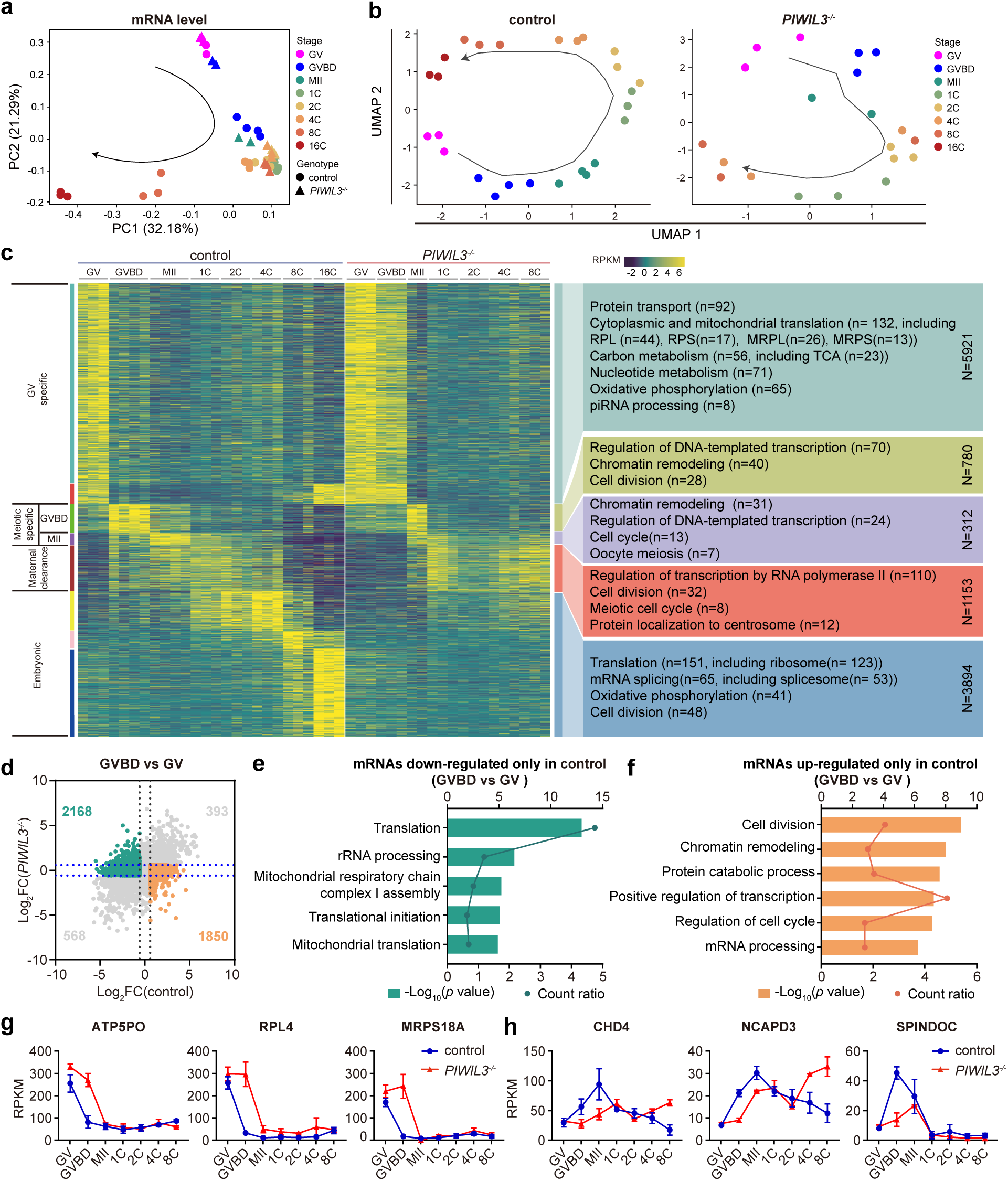
Impaired transcriptome reprogramming in *PIWIL3*-deficient oocytes and maternal *PIWIL3*-deficient embryos during the oocyte-to-embryo transition (OET). **a** PCA analysis of RNA levels expressed in the oocytes and embryos at various stages. Circles represent control (WT or *PIWIL3^+/-^*) oocytes and control (WT) embryos, while triangles represent *PIWIL3*-deficient (*PIWIL3^-/-^*) oocytes and *PIWIL3*-deficient (maternal *PIWIL3^-/-^*) embryos. **b** Pseudotime analysis of all gene expression using Monocle3 reveals distinct developmental trajectories in control (WT or *PIWIL3^+/-^*) and *PIWIL3*-deficient (*PIWIL3^-/-^*) oocytes and embryos. Arrowed curves represent simulated developmental sequences, demonstrating that *PIWIL3*-deficient samples exhibit disordered trajectories. **c** Heatmaps of differential mRNA expression modules in control and *PIWIL3*-deficient samples across developmental stages, normalized to RPKM. Each module includes the top 10% of genes with the highest eigengene connectivity (kME). Modules were grouped as GV oocyte-specific (turquoise, red), meiotic-specific (green (GVBD), purple (MII)), maternal clearance (brown), and embryonic-specific (yellow (minor ZGA), pink and blue (major ZGA)). Gene Ontology (GO) enrichment results are shown on the right; “n” denotes the number of enriched genes per pathway and “N” the total number of genes in each set. **d** Scatter plot showing log₂ fold change (FC) in control GVBD oocytes (x-axis) and *PIWIL3^-/-^*GVBD oocytes (y-axis) relative to GV oocytes. Each dot represents one gene; orange dots indicate genes significantly upregulated (log₂FC ≥ 1) only in control GVBD oocytes, while green dots indicate genes significantly downregulated (log₂FC ≤ -1) only in control GVBD oocytes. **e, f** GO enrichment analyses of the green-labeled (**e**) and orange-labeled (**f**) genes from panel d. The upper x-axis shows -log₁₀(*p* value) (bar length), and the lower x-axis shows the count ratio (dots and line). **g, h** Line plots showing representative genes from panel **d**. Green-labeled (**g**) genes significantly downregulated only in control GVBD oocytes. Orange-labeled (**h**) genes significantly upregulated only in control GVBD oocytes. Curves depict mRNA expression dynamics from the GV to 8C stages, with blue lines representing control samples and red lines representing *PIWIL3*-deficient samples.

By contrast, *PIWIL3*-deficient oocytes exhibited a near-complete failure of transcriptomic reprogramming at the GV-to-GVBD transition. Transcriptomes from mutant GVBD oocytes clustered with GV-stage profile but did not advance to the GVBD stage of controls (Fig. 3a). This defect persisted, as MII mutant oocytes retained transcriptomic features of control GVBD oocytes, indicative of severe developmental delay. Furthermore, maternal-*PIWIL3*-deficient 8C embryos clustered with control 4C embryos, reflecting developmental arrest at the 4C stage. Ultimately, no mutant embryos reached the 16C stage.

Pseudotime analysis revealed a highly disorganized trajectories in *PIWIL3*-deficient samples, in contrast to the continuous and orderly developmental trajectory from GV to 16C in controls (Fig. 3b). All of the above findings demonstrate that the loss of *PIWIL3* disrupts maternal mRNA regulation, leading to delayed oocyte maturation and persistent transcriptomic defects that impair embryonic progression.

To further dissect PIWIL3-dependent transcriptomic reprogramming, we applied weighted gene co-expression network analysis (WGCNA) across all control and mutant samples and identified eight distinct gene modules associated with PIWIL3 function: turquoise (5,386 genes), red (535 genes), green (780 genes), purple (312 genes), brown (1,253 genes), yellow (1,060 genes), pink (475 genes), and blue (2,359 genes) (Fig. 3c). Among these, the turquoise and red modules, specifically expressed at the GV stage, are hereafter referred to as *GV-specific* transcripts. These transcripts are normally silenced during the GV-to-GVBD transition, but remained expressed in *PIWIL3*-deficient oocytes up to the GVBD-to-MII transition. These genes encode components of cytoplasmic and mitochondrial ribosomes (RPL/RPS, MRPL/MRPS) as well as factors in protein transport, metabolism, oxidative phosphorylation, and piRNA processing. Thus, PIWIL3 is required for the timely silencing of GV-specific transcripts during meiotic resumption (Fig. 3c; Supplementary information, Fig. S3b).

Subsequently, three other oogenic modules (green, purple, and brown) undergo transcriptional activation during the GV-to-GVBD transition, complementing the widespread repression characteristic of this stage. In particular, the green and purple modules are strongly upregulated at GVBD and MII, respectively (Fig. 3c), represent *meiotic-specific* transcripts. In *PIWIL3*-deficient oocytes, the activation of these modules was markedly weakened and delayed. Functional enrichment revealed involvement in transcription, chromatin remodeling, and cell-cycle regulation, especially oocyte meiosis. PIWIL3 is therefore required for timely induction of meiotic regulators (Fig. 3c; Supplementary information, Fig. S3b).

The brown module corresponds to *maternal clearance* transcripts, a group of MZT-regulated maternal mRNAs that are normally activated around GVBD, maintained through MII, markedly down-regulated in zygote-4C embryos, and completely cleared by the 8C stage. In *PIWIL3*-deficient oocytes, however, these transcripts were not activated until MII, and failed to be down-regulated zygote-4C embryos, leading to their persistent expression through 8C, the latest stage we examined. Functionally, these genes are enriched for transcription by RNA polymerase II, cell division, and centrosome-related processes, indicating that PIWIL3 safeguards proper maternal clearance and the MZT (Fig. 3c).

Overall, PIWIL3 regulates activation of ∼2,345 genes (green, purple, brown modules) and repression of ∼5,921 genes (turquoise, red modules) around GVBD. Differential expression analysis (DESeq2) further confirmed that the GV-to-GVBD transition represents the most dynamic transcriptomic stage, with the largest number of differentially expressed genes (Supplementary information, Fig. S3a). In controls, 2,243 genes were upregulated and 2,736 downregulated at GVBD versus GV (FDR ≤ 0.05, fold change ≥ 2). However, in PIWIL3-deficient oocytes, only 393 of the 2,243 normally upregulated genes were properly activated, indicating a failure of transcriptional activation of 83% (1,850) of genes (Fig. 3d). Similarly, 80% (2,168) of the 2,736 normally repressed genes failed to be silenced in mutants (Fig. 3d). Gene Ontology analysis revealed that activation-defective genes were enriched in cell division, chromatin remodeling, protein catabolism, transcriptional regulation, and cell cycle, whereas repression-defective genes were enriched in translation, rRNA processing, mitochondrial respiratory chain assembly, translational initiation, and mitochondrial translation (Fig. 3e, f). Expression profiles of representative genes are shown in Fig. 3g, h. These observations revealed a key role of PIWIL3 in the most significant phase of transcriptomic programming during oocyte maturation.

In addition to modules associated with oocyte maturation, three other modules (yellow, pink, and blue) were activated during early embryogenesis, representing zygotically expressed *embryonic-specific* transcripts (Fig. 3c). Genes in the yellow module increased gradually from 1C to 4C, likely corresponding to minor ZGA, while pink and blue module genes were robustly induced after the 4C-to-8C transition, marking the onset of major ZGA in rabbits. In *PIWIL3*-deficient embryos, both minor and major ZGA gene cohorts failed to be activated, lacking transcriptional programs essential for embryogenesis. These genes are enriched in translation, mRNA splicing, oxidative phosphorylation, and cell division (Fig. 3c). Consistently, DESeq2 analysis revealed that 1,847 genes were activated during the major ZGA in controls, whereas only 11 in maternal-*PIWIL3*-deficient embryos (Supplementary information, Fig. S3a). These 1,847 genes encode factors critical for RNA processing, protein synthesis, and mitochondrial energy metabolism (Supplementary information, Fig. S3b). The failure of their activation provides a mechanistic explanation for the complete developmental arrest of mutants at the 8C stage.

In summary, *PIWIL3*-deficient samples diverged markedly from controls beginning at GVBD, with persistent massive misregulation throughout oocyte maturation and failure of ZGA during embryogenesis. PIWIL3 thus is a pivotal regulator of transcriptomic reprogramming during oocyte maturation and early embryogenesis in rabbits. This role contrasts sharply with golden hamsters, where PIWIL3 deficiency only affects the expression of 21 mRNAs in MII oocytes^30^.

### PIWIL3 is essential for proteomic reprogramming during the oocyte maturation and zygotic genome activation

The dynamic regulation of proteins, driven by precise protein synthesis, accumulation, and degradation, is pivotal for oocyte maturation, fertilization, and early embryogenesis ^3, 48–52^. To elucidate how PIWIL3 orchestrates proteomic dynamics during the OET, we performed high-resolution, single-cell proteomic profiling across developmental stages in both control and PIWIL3-deficient oocytes and embryos using liquid chromatography-tandem mass spectrometry (LC-MS/MS).

PCA of control proteomes revealed a structured developmental trajectory marked by three major transitions: GV to GVBD, GVBD to MII, and MII to the 1C (zygote) stage. In contrast, later embryonic stages (1C to 8C) exhibited limited separation (Fig. 4a). This was further supported by scatterplot analysis (Fig. 4b), which showed widespread upregulation of proteins during the GVBD-to-MII transition and subsequent downregulation from MII to the 1C stage—highlighting the unique proteomic landscape at MII.

**Fig. 4.**
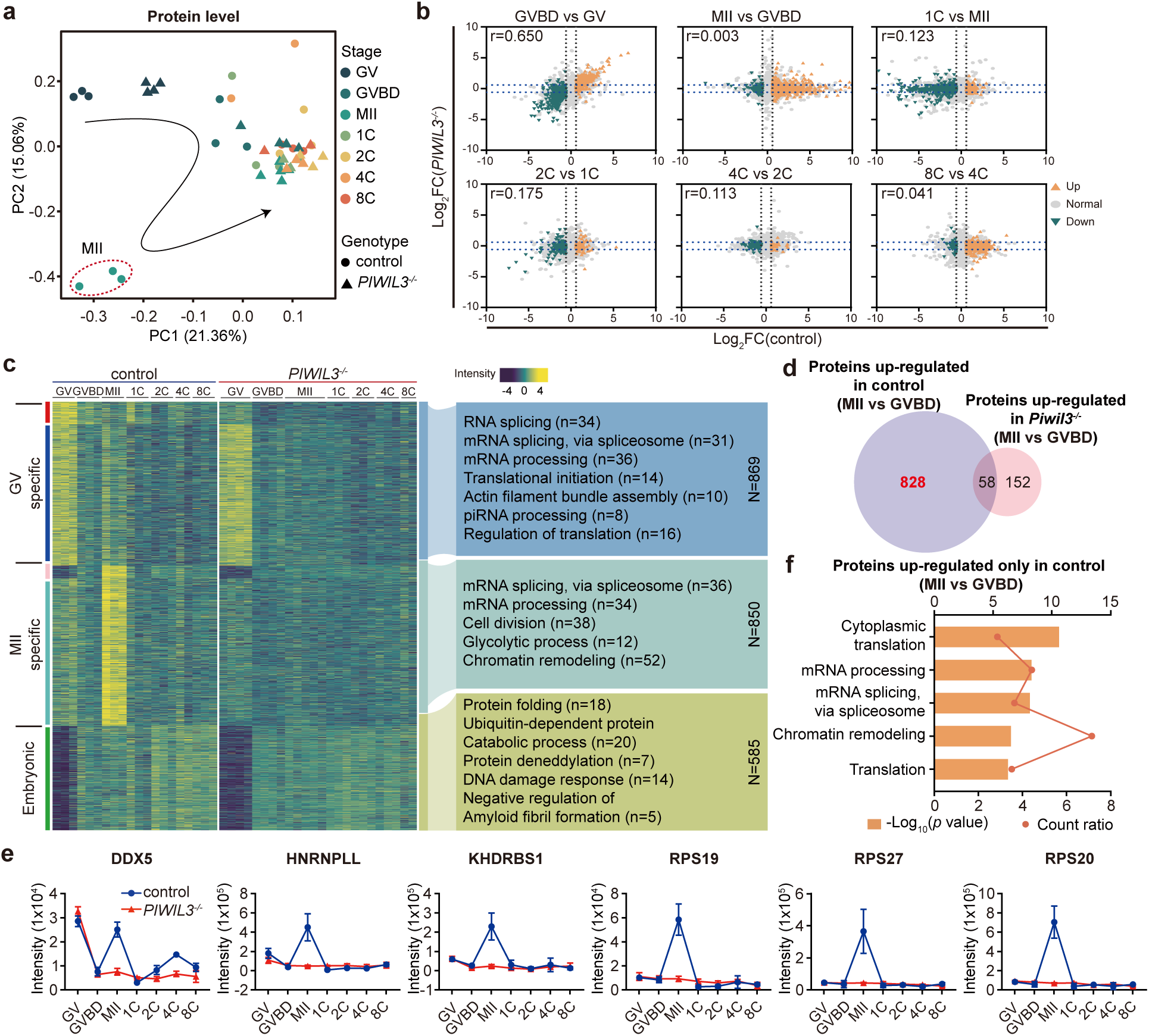
Impaired proteomic reprogramming in *PIWIL3*-deficient oocytes and maternal *PIWIL3*-deficient embryos during the OET. **a** PCA analysis of protein levels in the oocytes and embryos at various stages. Circles represent control (WT or *PIWIL3^+/-^*) oocytes and control (WT) embryos, while triangles represent *PIWIL3*-deficient (*PIWIL3^-/-^*) oocytes and *PIWIL3*-deficient (maternal *PIWIL3^-/-^*) embryos. **b** Scatter plots of differentially expressed proteins across successive stage transitions (GVBD versus GV, MII versus GVBD, 1C versus MII, 2C versus 1C, 4C versus 2C, 8C versus 4C) in control and *PIWIL3*-deficient samples. The x-axis shows log₂FC in control samples, and the y-axis shows log₂FC in *PIWIL3*-deficient samples. Significantly upregulated and downregulated proteins in control samples (≥ 2-fold, FDR ≤ 0.05) are highlighted in orange and green, respectively. r denotes the *Pearson* correlation coefficient between x and y values. **c** Heatmaps of differential protein expression modules in control and *PIWIL3*-deficient samples across developmental stages, normalized to protein intensity. Modules were grouped as GV oocyte-specific (red, blue), MII oocyte-specific (pink, turquoise), and embryonic-specific (green). GO enrichment results are shown on the right, with “n” indicating the number of genes per pathway and “N” the total number of genes in each set. **d** Venn diagram showing genes significantly upregulated in control (light purple) and *PIWIL3*-deficient (light pink) oocytes (fold change ≥ 2, *p* ≤ 0.05); genes detected only at MII but not GVBD oocytes were also included. **e** Line plots showing representative genes from panel f. Curves depict protein expression dynamics from the GV to 8C stages, with blue lines representing control samples and red lines representing *PIWIL3*-deficient samples. **f** GO enrichment analysis of the 828 genes uniquely upregulated in control oocytes. The upper x-axis shows -log₁₀(*p* value) (bar length), and the lower x-axis shows the count ratio (dots and line).

By contrast, *PIWIL3*-deficient samples showed an aberrant proteomic trajectory. First, the mutant GV oocytes show a distinct profile between the control GV and GVBD. Second, the mutant GVDB, MII, 1C-8C embryos all collapsed to one cluster (Fig. 4A, 4B), indicating a failure to undergo the critical proteomic shift starting at the GVBD-MII transition. These findings demonstrate that PIWIL3 is indispensable for proper proteomic reprogramming during the OET, particularly around the MII stage.

To further resolve the temporal dynamics of the proteome during OET, we identified four PIWIL3-regulated protein co-expression modules: Red (120 proteins), Blue (749 proteins), Turquoise (782 proteins), and Pink (68 proteins) (Fig. 4c). Both Red and Blue modules showed high protein expression specifically at the GV stage, but were sharply down-regulated to basal levels at the GVBD stage, and remained so up to 8C stage, with blue module proteins partially re-accumulating at MII. GO analysis indicated that these modules are enriched for RNA splicing, mRNA processing, translational initiation, actin filament bundle assembly, and piRNA processing. In contrast, the pink and turquoise modules peaked at MII and were associated with mRNA splicing, glycolysis, cell cycle regulation, and cell division (Fig. 4c). Collectively, these four modules accounted for nearly 30% of the total proteome, *PIWIL3* deficiency abolished the expression of most of the above proteins in all four modules, except for the GV-specific expression of blue module proteins, during the entire course of OET (Fig. 4c). This indicates that PIWIL3 regulates the expression of a broad spectrum of proteins during oocyte maturation.

Supporting this, during the GVBD-to-MII transition, 886 proteins in control oocytes showed robust accumulation at MII (Fig. 4d), including 464 that were significantly upregulated (fold change ≥ 2, *p* ≤ 0.05) and 422 detected exclusively at MII (Supplementary Fig. S4c). In contrast, in *PIWIL3*-deficient oocytes, only 58 of the 828 proteins retained high expression (Fig. 4d), with the representative protein expression profiles shown in Fig. 4e.

GO analyses revealed that these PIWIL3-controlled proteins are primarily involved in cytoplasmic translation, mRNA processing, mRNA splicing via the spliceosome, and chromatin remodeling (Fig. 4f). These data indicate that PIWIL3 is a key regulator of stage-specific proteomic remodeling during oocyte maturation, especially for the substantial protein accumulation at MII and subsequent ZGA (also see below).

In addition to oogenic modules, we identified a green module whose proteins increased significantly after the 1C stage, coinciding with the onset of minor ZGA in early embryogenesis. These proteins are likely translated from maternal mRNAs to facilitate ZGA, and maternal *PIWIL3* deficiency disrupted their expression dynamics from the 1C to 8C stages (Fig. 4c). Differential protein expression analysis further confirmed that PIWIL3 is required for ZGA (Supplementary information, Fig. S4a-c). In summary, our findings establish PIWIL3 as a master regulator of proteomic reprogramming during both oocyte maturation and ZGA.

### PIWIL3 directly regulates target transcripts at RNA and protein levels during the OET

To investigate how PIWIL3 regulates transcriptomic and proteomic reprogramming during the OET, we performed RNA immunoprecipitation followed by mRNA sequencing (PIWIL3-RIP-mRNAseq) on GV to GVBD oocytes to identify direct mRNA targets of PIWIL3. 1,060 mRNAs were bound by PIWIL3 across biological replicates (Fig. 5a). KEGG analysis of these mRNAs revealed significant enrichment in pathways related to the spliceosome, cell cycle, mRNA surveillance, ribosome biogenesis, oxidative phosphorylation, cellular senescence, and oocyte meiosis (Fig. 5b).

**Fig. 5.**
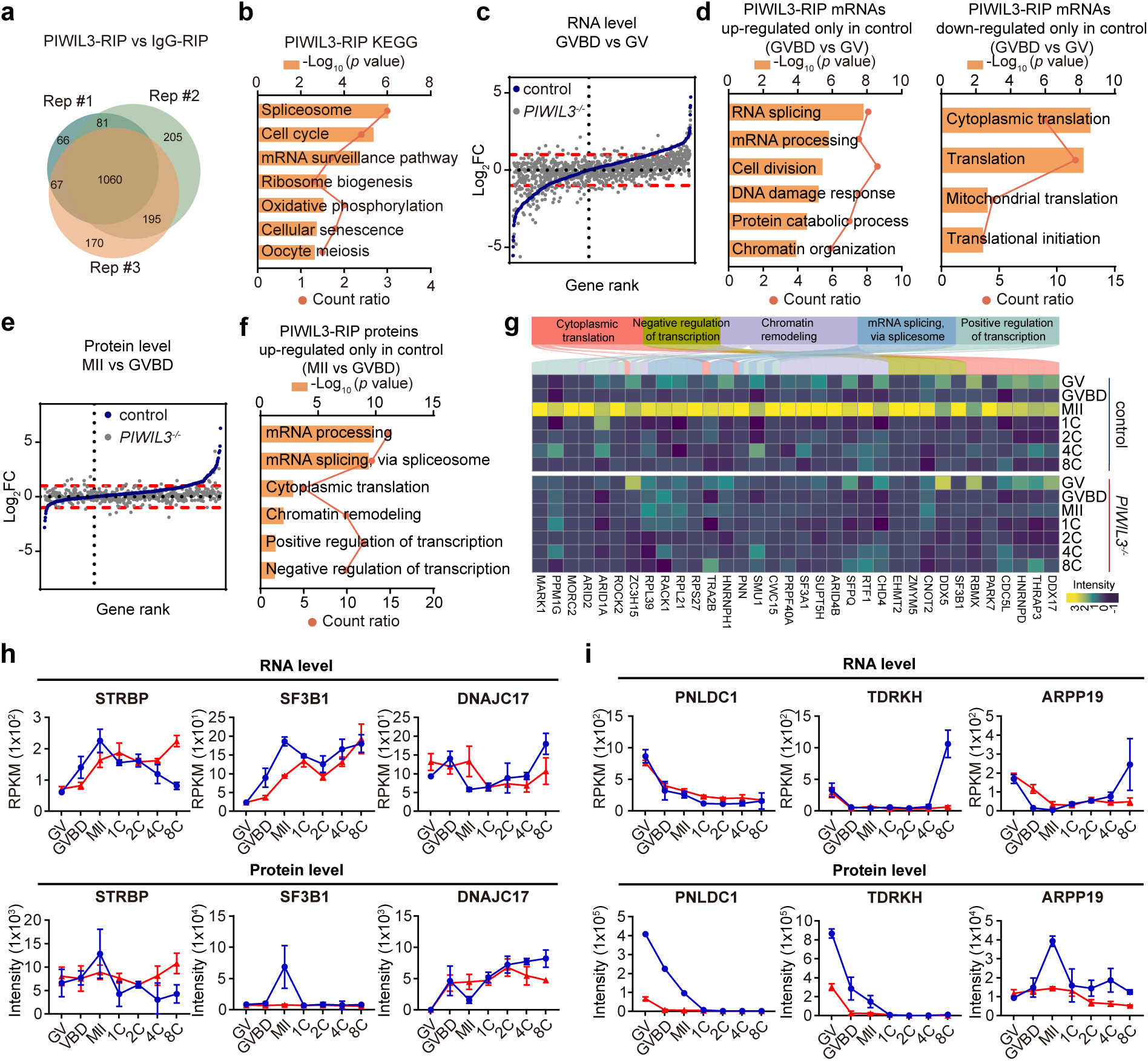
PIWIL3 directly targets transcripts at the RNA and Protein levels. **a** Venn diagram showing the overlap of 1,060 mRNAs bound by PIWIL3 across three independent PIWIL3-RIP-mRNAseq experiments (Rep_#1, Rep_#2, and Rep_#3). **b** KEGG pathway analysis of 1,060 mRNAs bound by PIWIL3. **c** Scatter plot of PIWIL3-bound mRNAs (from panel a) showing RNA expression changes during the GV-GVBD transition. Each point represents one gene, ranked along the x-axis by the log₂FC in control oocytes from lowest to highest (gene rank). The y-axis indicates the log₂FC of GVBD versus GV expression. Dark blue points represent PIWIL3-bound mRNAs in control oocytes, and grey points represent the same mRNAs in PIWIL3-deficient oocytes. Red dashed lines mark log₂FC = 1 and -1 thresholds, and the vertical grey dashed line marks the x-axis position corresponding to log₂FC = 0. **d** GO enrichment analysis of PIWIL3-bound mRNAs from panel **c** that were significantly upregulated (log₂FC ≥ 1, left) or downregulated (log₂FC ≤ -1, right) during the GV-GVBD transition only in control oocytes. The upper x-axis shows -log₁₀(*p* value) (bar length), and the lower x-axis shows the count ratio (dots and line). **e** Scatter plot of PIWIL3-bound mRNAs (from panel a) showing protein expression changes during the GVBD-MII transition. Each point represents one gene, ranked along the x-axis by the log₂ fold change (FC) in control oocytes from lowest to highest (gene rank). The y-axis indicates the log₂FC of MII versus GVBD expression. Dark blue points represent PIWIL3-bound mRNAs in control oocytes, and grey points represent the same mRNAs in PIWIL3-deficient oocytes. Red dashed lines mark log₂FC = 1 and -1 thresholds, and the vertical grey dashed line marks the x-axis position corresponding to log₂FC = 0. **f** GO enrichment analysis of PIWIL3-bound mRNAs from panel **e** that were significantly upregulated (log₂FC ≥ 1) during the GVBD–MII transition only in control oocytes. The upper x-axis shows -log₁₀(*p* value) (bar length), and the lower x-axis shows the count ratio (dots and line). **g** Integrated heatmap and Sankey plot showing protein expression dynamics of the clustered genes across GV to 8C stages in control and *PIWIL3*-deficient samples. Genes are grouped by their GO term assignments (from panel **f**), with the corresponding pathways indicated by colored boxes above the heatmap. **h** Genes in which protein regulation by PIWIL3 paralleled mRNA dynamics. **i** Genes in which protein regulation by PIWIL3 was uncoupled from transcript levels. blue lines indicate control samples and red lines *PIWIL3*-deficient samples.

We then examined the dynamics of these transcripts and their protein products during oocyte maturation. As the GV-to-GVBD transition represents the most dynamic window of transcriptomic reprogramming (Fig. 3a-d), we focused on this stage to assess direct effects of PIWIL3. In control oocytes, numerous PIWIL3-bound mRNAs showed marked changes between GV and GVBD (Fig. 5c; Supplementary information, Fig. S5a). Transcripts associated with cell division, RNA splicing, mRNA processing, and chromatin organization were significantly upregulated, whereas those related to translation were downregulated (Supplementary information, Fig. S5b-c). In contrast, in *PIWIL3*-deficient oocytes, ∼85% of the PIWIL3-bound mRNAs failed to show significant changes (Fig. 5c; Supplementary information, Fig. S5a). GO analyses indicated that transcripts failing to increase in mutants were enriched for cell division, RNA processing, and chromatin organization, whereas those failing to be degraded were mainly associated with cytoplasmic and mitochondrial translation (Fig. 5d). These results demonstrate that, PIWIL3 directly regulates numerous mRNAs that act upstream in regulatory hierarchies during the GV-to-GVBD transition, positioning PIWIL3 as a key regulator of the post-transcriptional networks that orchestrate transcriptomic reprogramming essential for oocyte development.

Since *PIWIL3* deficiency blocks the extensive protein accumulation in the MII oocytes (Fig. 4a-d; Supplementary information, Fig. S4a-b), we investigated to what extent the blockage was due to direct PIWIL3 regulation by analyzing proteins encoded by PIWIL3-bound mRNAs. In controls, these proteins were generally upregulated from GVBD to MII (Fig. 5e). In mutants, however, levels of these proteins remained largely unchanged, including splicing factors, transcription factors, translation machinery, and chromatin remodelers (Fig. 5f, g), indicating that PIWIL3 directly controls the proteomic remodeling of many gene expression regulators during oocyte maturation.

Mechanistically, PIWIL3 appears to regulate protein abundance through multiple layers, including control of mRNA stability, translational efficiency, and post-translational stability. In some cases, PIWIL3-dependent defects in protein expression mirrored altered mRNA dynamics (Fig. 5h), whereas in others protein abundance was uncoupled from transcript levels (Fig. 5i), suggesting additional regulation at translational or post-translational levels.

### PIWIL3 is essential for piRNA biogenesis in rabbit oocytes and embryos

Having observed transcriptomic, proteomic, and phenotypic defects resulting from PIWIL3 deficiency, we sought to determine how PIWIL3 exerts its regulatory functions. PIWI proteins typically partner with piRNA to regulate gene expression at multiple levels. To test if this is also the case for PIWIL3, we conducted single-cell small RNA sequencing to profile piRNA dynamics across all stages of oocyte maturation and early embryogenesis (from GV to 8C) and assessed the impact of PIWIL3 deficiency.

In control samples, piRNAs constituted the vast majority of small RNAs in both oocytes and early embryos, with a predominant population peaked at 18 nucleotides (nt; Fig. 6a). The abundance of 18-nt piRNAs, as normalized to spike-in sequences, was relatively low in GV and GVBD oocytes but increased dramatically as oocytes matured to the MII stage and remained so in 1C embryos. From the 2C to 8C stages, piRNA abundance fluctuated noticeably but remained high overall, dominating the small RNA landscape as seen in oocytes (Fig. 6a).

**Fig. 6.**
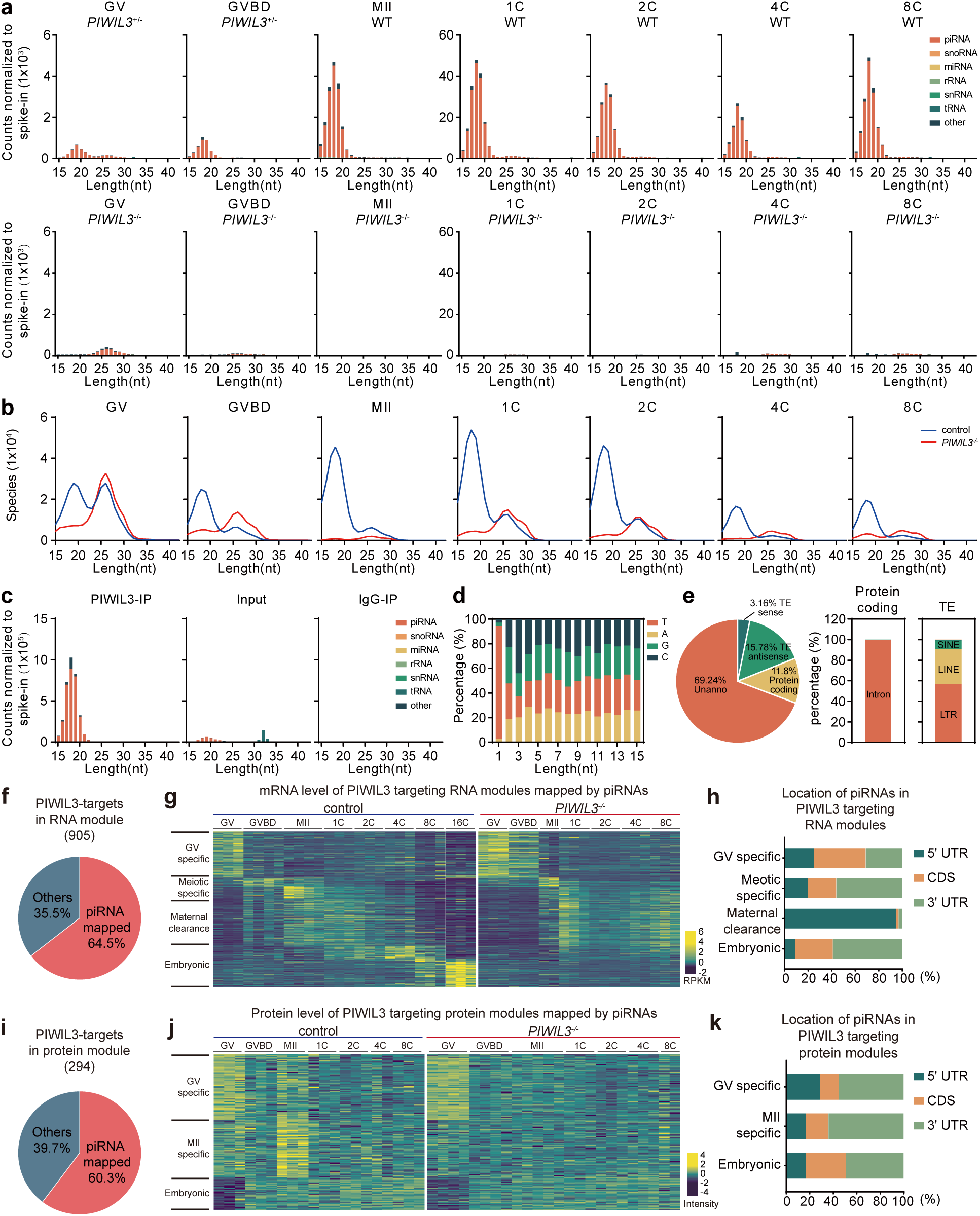
Characterization of small RNAs in *PIWIL3*-mutant rabbits. **a** Histogram illustrating composition of small RNA categories by length distribution. Small-RNA reads were classified into tRNAs, rRNAs, small nuclear RNAs (snRNAs), small nucleolar RNAs (snoRNAs), microRNAs (miRNAs), and piRNA candidates, each represented by distinct colors. Small RNA counts were normalized to the exogenous spike-in. Data represent the average value of biological replicates at each time point: n = 3 (GV), n = 4 (GVBD), n = 3 (MII), n = 5 (1C), n = 5 (2C), n = 4 (4C), n = 5 (8C) for control (WT or *PIWIL3^+/-^*) ; n = 5 (GV), n = 4 (GVBD), n = 3 (MII), n = 4 (1C), n = 4 (2C), n = 4 (4C), n = 4 (8C) for *PIWIL3*-deficient (*PIWIL3^-/-^*) samples. **b** Curve graph showing the size distribution of unique piRNA sequences in control (blue) and *PIWIL3*-deficient samples (red). **c** Histogram showing the size distribution of PIWIL3-bound piRNAs. PIWIL3-bound piRNAs were identified by immunoprecipitation using anti-PIWIL3 antibodies from WT GV and GVBD oocytes. Mouse non-specific immunoglobulin G (IgG) antibodies served as a negative control. The bound piRNA population consisted of sequences ranging from 16 to 20 nucleotides in length. **d** Bar graph depicting the nucleotide preference profile of PIWIL3-bound piRNAs. **e** Pie chart (left) showing the genome annotation of PIWIL3-bound piRNAs, bar graph depicting the ratio of exons to introns within protein coding genes (middle) and various types within TE families (right). **f** Pie chart showing mapping of PIWIL3-bound piRNAs to 905 PIWIL3-bound mRNAs identified in PIWIL3-regulated RNA modules (Fig. 3c). Mapping was performed with a 15-nt seed match and zero mismatches; 64.5% of these transcripts were mapped and defined as PIWIL3 direct targets. **g** Heatmap of the mapped PIWIL3 direct targets from panel f showing mRNA dynamics from GV to 8C in control and PIWIL3-deficient (*PIWIL3^-/-^*) samples, normalized to RPKM. Targets were grouped into GV-specific, meiotic-specific, maternal clearance, and embryonic categories according to their stage-specific expression patterns. **h** Distribution of PIWIL3-bound piRNA target sites within the transcript groups in panel **g**. Bars indicate the percentage of sites mapping to the 5′UTR, CDS, or 3′UTR. **i** Pie chart showing mapping of PIWIL3-bound piRNAs to 294 PIWIL3-bound mRNAs identified in PIWIL3-regulated protein modules (Fig. 4c). Mapping was performed with a 15-nt seed match and zero mismatches; 60.3% of these transcripts were mapped and defined as PIWIL3 direct targets. **j** Heatmap of the mapped PIWIL3 direct targets from panel **f** showing protein dynamics from GV to 8C in control and PIWIL3-deficient (*PIWIL3^-/-^*) samples, normalized to intensity. Targets were grouped into GV-specific, meiotic-specific, maternal clearance, and embryonic categories according to their stage-specific expression patterns. **k** Distribution of PIWIL3-bound piRNA target sites within the transcript groups in panel **j**. Bars indicate the percentage of sites mapping to the 5′UTR, CDS, or 3′UTR.

PIWIL3 mutations completely abolished the biogenesis of 18 nt piRNA population. In *PIWIL3*-deficient GV oocytes, a low abundant population of small RNAs peaking at 26 nt appeared (Fig. 6a). In GVBD oocytes, the 26-nt population was significantly reduced. By the MII stage, 26-nt piRNAs were also undetectable (Fig. 6a). In 1C-8C mutant embryos, piRNAs remained undetectable (Fig. 6a). These results indicate that PIWIL3 is essential for piRNA biogenesis in oocytes and early embryos.

We then analyzed what piRNA species are produced at different stages of OET and how they are affected by PIWIL3 deficiency. Among control samples, GV oocytes contained equally complex 18-nt and 26-nt piRNA populations, although the abundance of the 18-nt piRNA population was much higher than that of the 26-nt piRNA population (Fig. 6b). In GVBD oocytes, the complexity of the 18-nt population was maintained, while that of the 26-nt population drastically decreased. By the MII stage, the complexity of 18-nt piRNAs doubled, whereas the 26-nt population became nearly undetectable (Fig. 6b). These findings reveal a shift to 18-nt piRNA biogenesis as oocytes mature. These 18-nt piRNA persisted as the dominant population throughout early embryonic stages (1C to 8C). Upon zygotic genome activation around the 4C stage, the diversity of both 18-nt and 26-nt piRNA populations significantly declined.

In contrast, *PIWIL3*-deficient oocytes and embryos contained mostly the 26-nt unique RNAs, with 18-nt population nearly absent (Fig. 6b). Specifically, mutant GV oocytes contained few species of 18-nt piRNAs but a complex population of the 26-nt piRNAs. As oocytes matured to the GVBD stage, the complexity of 26-nt piRNA significantly decreased. Eventually, few species of piRNAs were detected in MII oocytes (Fig. 6b). In maternal *PIWIL3*-deficient embryos, the 18-nt piRNA species continued to be absent, and the complexity of the 26-nt population gradually declined, paralleling trends observed in control embryos (Fig. 6b). Collectively, these findings confirms that PIWIL3 is essential for piRNA biogenesis and acts as a critical regulator of the entire small RNA landscape during rabbit oogenesis and early embryogenesis.

### PIWIL3 binds to the 18-nt piRNA population enriched in 5’U and mostly encoded by unannotated regions

We then investigated whether PIWIL3-bound piRNAs correspond to the abundant 18-nt population and whether PIWIL3 forms complexes with these piRNAs to mediate gene regulation. We sequenced PIWIL3-bound small RNAs from rabbit GV and GVBD oocytes and found that PIWIL3-bound piRNAs predominantly consisted of 18-nt species (Fig. 6c). These piRNAs exhibited a strong 5’ uridine bias (Fig. 6d) but lacked significant adenine enrichment at position 10. This nucleotide signature indicates, together with the predominat presence of PIWIL3 in oocytes and early embryos, indicate that PIWIL3-bound piRNAs are likely generated through a primary piRNA biogenesis pathway.

To further understand the piRNA biogenesis during oogenesis and early embryogenesis, we examined the expression of other canonical piRNA biogenesis factors. MAEL, ASZ1, TDRKH, TDRD1, TDRD5, DDX4, RNF17, PNLDC1, and HENMT1 were prominently expressed in GV oocytes, in addition to PIWI proteins (PIWIL1, PIWIL2, PIWIL4) (Supplementary information, Fig. S6a, b). Notably, PIWIL3 deficiency reduced protein levels of several core factors, including ASZ1, TDRKH, PNLDC1, HENMT1 and GTSF1 (Supplementary information, Fig. S6b). This effect likely contributes to the defect in piRNA biogenesis. In addition, both PIWIL1 mRNA and protein exhibited a higher expression in mutants compared to controls. This compensatory effect may account for the elevated abundance of 26-nt piRNAs in *PIWIL3*-deficient GV oocytes. This effect diminished beyond the GV stage, as PIWIL1 expression was reduced in both control and mutant oocytes after GV stage.

Genomic mapping revealed that ∼70% of PIWIL3-bound piRNAs originated from unannotated regions. Within the remaining sequences, 15.8% aligned antisense and 3.2% sense, respectively, to transposable elements (TEs). These TEs include long terminal repeats (LTRs, 65%), LINEs (28%) and SINEs (7%). In addition, 11.8% of PIWIL3-bound piRNAs mapped to protein-coding genes (Fig. 6e), markedly higher than in golden hamsters (3.5%)^30^. These data imply PIWIL3-piRNA complexes in rabbits may directly regulate the expression of mRNAs during oogenesis and early embryogenesis.

### PIWIL3 regulation of target mRNA appears to involve piRNA

We next asked whether PIWIL3-associated piRNAs contribute to regulation of target mRNAs. Among 905 PIWIL3-bound transcripts dynamically regulated during the OET, 64.5% contained perfect seed matches to PIWIL3-bound piRNAs within their 15-nt seed regions (Fig. 6f). These putative piRNA-targeted transcripts displayed stage-specific regulation: GV-specific mRNAs were negatively regulated during the GV-to-GVBD transition; meiotic-specific mRNAs were positively regulated at GVBD; MZT-associated maternal clearance transcripts were upregulated during oocyte maturation but downregulated at ZGA; and embryonic-specific transcripts were strongly induced during minor and major ZGA (Fig. 6g).

At the protein level, 294 PIWIL3-regulated targets were identified, 60.3% of which are translated from mRNAs harboring perfect 15-nt seed complementarity to PIWIL3-bound piRNAs (Fig. 6i). Maternal transcripts were mainly regulated at MII, whereas embryonic-specific targets exhibited a marked increase at the 2C stage following minor ZGA (Fig. 6j).

Overall, these findings are consistent with a major role of PIWL3 in regulating oogenesis and early embryogenesis, which sharply contrast with hamster oocytes, where PIWIL3 minimally influences mRNA regulation and PIWIL1-bound piRNAs do not directly target mRNAs ^30, 45^.

We then compared piRNA targeting patterns between transcripts altered at the RNA versus protein level. Among PIWIL3-target mRNAs with dynamic transcript changes, putative piRNA binding sites were distributed across 5’ UTRs, CDS, and 3’ UTRs in GV-specific transcripts that were negatively regulated during GVBD. However, in meiotic-specific transcripts positively regulated by PIWIL3, putative piRNA binding sites were strongly enriched in 3’ UTRs (56%). In MZT-associated maternal clearance transcripts, nearly all sites (95%) were localized to 5’ UTRs, whereas in embryonic-specific transcripts induced during ZGA, 3’ UTRs predominated (59%) (Fig. 6h). These distinct localization pattern might be indicative of different modes of regulation, since mRNA stability regulation can be achieved throughout the entire length of mRNA while translational regulation are known to involve 5’ UTR and 3’ UTR only.

Consistent with the above hypothesis, in mRNAs showing protein-level regulation, piRNA target sites were consistently enriched in 3’ UTRs (49–69%) across stages, with fewer sites in CDS (16–34%) and 5’ UTRs (17–29%) (Fig. 6k). These dynamic, stage-specific targeting patterns implicates that PIWIL3-piRNA complexes can achieve post-transcriptional regulation at multiple levels.

### PIWIL3 plays both suppressing and activating roles in regulating the stage-specific expression of transposable elements in rabbit oocytes and early embryos

PIWI proteins are widely known for their role in silencing transposable elements (TEs), such as LINEs, SINEs, and LTRs that serve as sources of piRNAs and regulators of chromatin and gene expression. While TE regulation by PIWI–piRNA complexes is well documented, their function in the female germline of mammals remains poorly understood. In golden hamsters, loss of PIWIL1 results in a over twofold upregulation of 29 TE families, whereas PIWIL3 deficiency has minimal effect on TE expression in oocytes and embryos^30, 31^. To investigate whether PIWIL3 regulates TEs in rabbit oocytes and early embryos, we analyzed SMART-seq3 data spanning from GV oocytes to 8C embryos from both *PIWIL3*-deficient and control females; 16C control embryos were also included to capture post-ZGA stages.

PCA of TE expression profiles revealed a clear stage-dependent clustering from GV oocytes to 16C embryos in control samples (Fig. 7a). GV oocytes, GVBD-4C stages, 8C embryos, and 16C embryos emerge as four distinct clusters. In contrast, *PIWIL3*-deficient samples exhibited a disrupted progression: GV and GVBD oocytes clustered with control GV oocytes, while MII oocytes and embryos from 1C to 8C stages clustered tightly with the GVBD-4C cluster in controls (Fig. 7a). This disruption suggests that PIWIL3 ensures the proper progression of stage-specific expression of TEs during oocyte maturation and early embryogenesis.

**Fig. 7.**
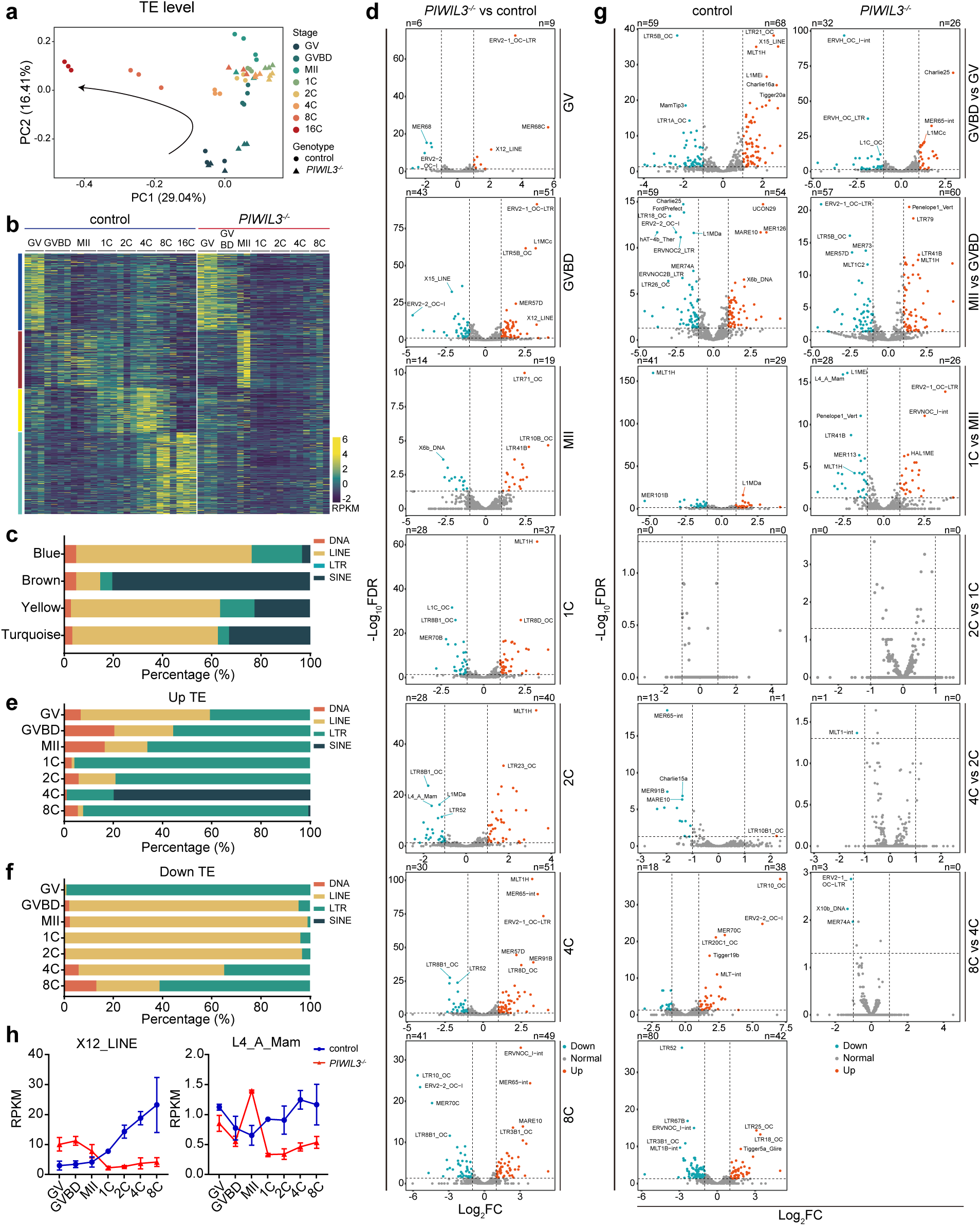
Altered transposable element (TE) expression in *PIWIL3*-deficient oocytes and embryos during the OET. **a** PCA analysis of TE families expressed in the oocytes and embryos at various stages. Circles represent control (WT or *PIWIL3^+/-^*) oocytes and control (WT) embryos, while triangles represent *PIWIL3*-deficient (*PIWIL3^-/-^*) oocytes and *PIWIL3*-deficient (maternal *PIWIL3^-/-^*) embryos. **b** Heatmap showing differential TE expression patterns of TE modules in control and *PIWIL3*-deficient samples across developmental stages, normalized to RPKM. **c** Bar graph depicting the types and distribution proportions of TEs within the four modules. **d** Volcano plots depicting differentially expressed TEs between control and *PIWIL3*-deficient samples (≥ 2-fold change; FDR ≤ 0.05), Highly significant upregulated and downregulated TEs are highlighted in orange and green, respectively. **e-f** Bar graph depicting the types and distribution proportions of significant upregulated (**e**) and downregulated (**f**) TEs between control and *PIWIL3*-deficient samples across various developmental stages. **g** Volcano plots illustrating differentially expressed TEs across the following stage comparisons in both control and *PIWIL3*-deficient samples: GVBD versus GV; MII versus GVBD; 1C versus MII; 2C versus 1C; 4C versus 2C; and 8C versus 4C. Significantly upregulated and downregulated TEs (≥ 2-fold change; FDR ≤ 0.05) are highlighted in orange and green, respectively. **h** Line plots illustrating dynamic TE expression changes of representative TEs during the OET (GV to 8C) in control (blue) and *PIWIL3*-deficient (red) samples.

To further characterize PIWIL3 regulation of TE expression, we performed WGCNA, which identified four distinct TE expression modules (Fig. 7b). Among them, the Blue and Brown modules were predominantly active during oogenesis. The Blue module, primarily composed of LINE and LTR elements (with LINEs being dominant; Fig. 7c), was specifically active in control GV oocytes, but showed expanded activity at both GV and GVBD stages in PIWIL3-deficient oocytes. The Brown module, enriched for SINE elements (Fig. 7c), displayed progressive expression from GV to MII stages in control oocytes and remained at low levels during the 1C–4C stages. However, in *PIWIL3*-deficient oocytes, this module was abnormally restricted to the MII stage at which it was expressed at significantly higher levels than controls (Fig. 7b). These findings indicate that PIWIL3 primarily functions to suppress premature or excessive activation of LINE, LTR, and SINE elements during oocyte maturation.

Two additional modules, Yellow and Turquoise, were activated post-fertilization in control embryos, mostly at 2C-16C stages, coinciding with the onset of ZGA. The Yellow module, consisting primarily of other LINEs, SINEs and LTRs, showed a marked increase in expression from the 2C to 4C stages in control embryos (Fig. 7b, c). The Turquoise module, dominated by a third set of LINE elements, exhibited low expression at the 4C stage and peaked at the 8C and 16C stages (Fig. 7b, c). In maternally *PIWIL3*-deficient embryos, however, neither the yellow nor turquoise modules were activated from 1C to 8C (Fig. 7b), suggesting that PIWIL3 is required for activation of ZGA-associated LINE, LTR, and SINE elements, which differ from those expressed during oogenesis.

Analysis of individual TE families further confirmed the dual regulatory roles of PIWIL3 (Fig. 7d, g). For example, X12_LINE, an embryo-specific TE, was abnormally expressed in *PIWIL3*-deficient oocytes but failed to be activated during embryogenesis (Fig. 7h). L4_A_Man, normally expressed at a low level in MII oocytes but highly expressed in embryos, become highly expressed in *PIWIL3*-deficient MII oocytes and fail to achieve high expression during embryogenesis (Fig. 7h). These findings demonstrate that PIWIL3 serves both repressive and activating functions in TE regulation, depending on the developmental context—a function that contrasts with the repressive-only role typically observed for PIWI proteins in the male germline.

Subtype-specific analysis of deregulated TEs revealed that LTR elements were the most frequently upregulated upon loss of PIWIL3, particularly during embryogenesis (Fig. 7d-e). An exception was observed at the 4C stage, where upregulated TEs were primarily SINE elements (Fig. 7e). Interestingly, LINE elements constituted the majority of downregulated TEs from GVBD through the 4C stage, while LTRs were the dominant downregulated subtype at the GV and 8C stages (Fig. 7e-f). The near-exclusive presence of LTRs among downregulated TEs at the GV stage further emphasizes the stage-specific and subtype-selective regulatory function of PIWIL3.

Together, these results establish that PIWIL3 exerts dynamic, and stage-specific control over TE expression during rabbit oocyte maturation and early embryonic development, functioning as both a repressor and activator. This dual regulatory capacity highlights a unique role for PIWIL3 in the female germline.

### PIWIL3 regulation of transposons also appears to involve piRNA

Given the known association between PIWI proteins and piRNAs, we next examined whether PIWIL3-bound piRNAs were involved in regulating the differentially expressed TEs. Among all dysregulated TEs in *PIWIL3*-deficient samples, 67.4% harbored sequences with single-mismatch complementarity to PIWIL3-bound piRNAs within their 15-nt seed regions (Supplementary information, Fig. S7a). Both sense and antisense PIWIL3-bound piRNAs were broadly distributed across TEs that were either up- or downregulated (Supplementary information, Fig. S7b–d, S8). Interestingly, PIWIL3-bound piRNAs were more enriched in TEs that were downregulated in *PIWIL3*-deficient MII oocytes than in those that were upregulated (Supplementary information, Fig. S7d), suggesting a potential activating role for some piRNAs, depending on stage and TE subtype. This finding diverges from the classical model of piRNAs acting solely as repressors and stands in contrast to the minimal TE regulatory role of PIWIL3 observed in golden hamster oocytes. These findings are consistent with the involvement of piRNA in PIWIL3 regulation of TEs.

## DISCUSSION

### PIWIL3 is essential for oogenesis in higher mammals

The role of PIWI proteins in mammalian female reproduction has long been overlooked because most investigations used mice, in which *PIWIL3* does not exist yet gene knockouts of other *PIWI* genes only impact male fertility. This led to a long-time assumption that the *PIWI*-piRNA pathway is involved in female fertility in mammals. However, recent studies of *PIWIL3* in golden hamsters, another rodent species, demonstrate that PIWIL1 and PIWIL3 are expressed at high levels in oocytes, with PIWIL1 expression being predominant^29, 31^. Disrupting *PIWIL1* or *PIWIL3* in golden hamster causes females infertility and subfertility, respectively^30–32, 45^, indicating an essential role of PIWIL1 but a minor role of PIWL3 in female fertility. Of note, both proteins are not required for oogenesis, but only functions in early embryogenesis. Our work in rabbits, phylogenetically more advanced than rodents, demonstrates that, among the four PIWI genes, only *PIWIL3* is expressed in GVBD/MII oocytes and is essential for oogenesis. Specifically, *PIWIL3*-deficient female rabbits exhibit severe reductions in primary, secondary, and antral follicles. This contrasts with PIWIL3 in golden hamsters, where its deficiency does not lead to detectable defects in oocyte reserve ^30, 45^.

### PIWIL3 is essential for embryogenesis in higher mammals

Our work also reveals the essential function of maternal PIWIL3 for early embryonic mitotic divisions. Embryos lacking maternal PIWIL3 fail to undergo synchronous divisions and arrest by the 8-cell stage. This also contrasts with golden hamsters, where maternal PIWIL3 deficiency leads to only partial developmental arrest, and approximately 40% of embryos are still able to progress to the morula and blastocyst stages ^30, 45^. Importantly, rabbit PIWIL3 is more homologous to its human counterpart than to most other mammalian PIWIL3, second only to rhesus monkeys (Supplementary information, Fig. S1a).

### PIWIL3 binds to 18-nt piRNAs and is required for piRNA biogenesis

Both rabbit and human oocytes predominantly contain short piRNAs bound to PIWIL3^50^ (Fig. 6a, b). We have shown that PIWIL3 exclusively binds to 18-nt piRNAs, which is the predominate population of piRNA just like in humans. Furthermore, the loss of PIWL3 leads to failure of piRNA biogenesis from the GVBD stage onward. In contrast, golden hamster oocytes primarily harbor 23- and 29-nt piRNAs associated with PIWIL1^30–32^. PIWIL3 depletion in golden hamster does not alter either total piRNA abundance or complexity, possibly due to compensatory upregulation of PIWIL1-associated piRNAs^30, 45^. Thus, the rabbits is a better model system for studying human PIWI-piRNA biology. Our findings provide a framework to reinterpret unexplained human infertility cases—subjects with oocyte maturation defects or embryonic arrest may harbor undiagnosed mutations in PIWIL3 or associated piRNA factors, overlooked due to rodent-centric paradigms.

### PIWIL3 achieves its oogenic and embryogenic functions by regulating a large number of mRNAs via distinct mechanisms

Our study reveals that PIWIL3 regulates the expression of a large number of genes during oogenesis and early embryogenesis at both the mRNA and protein levels. Multi-omics analyses revealed 905 RNA-level and 294 protein-level targets under direct PIWIL3 control, spanning cell division, metabolism, chromatin remodeling, and translational regulation (Fig. 6f, g, i, j). In contrast, PIWIL3 in golden hamsters plays no obvious role in regulating gene expression during oocyte maturation and early embryogenesis ^30, 45^. Our findings establish PIWIL3 as a central post-transcriptional regulator of gene expression for its essential roles in oogenesis and early embryogenesis.

Notably, transcript-level regulation involves stage-specific piRNA localization: piRNAs map predominantly to CDS in GV oocytes, to 3’ UTRs during meiosis, to 5’ UTRs during MZT-associated maternal clearance, and again to 3’ UTRs during embryonic stages (Fig. 6g-h). In contrast, protein-level targets consistently exhibit piRNA binding mainly in 3’ UTRs across all stages (Fig. 6j-k), implying distinct mechanisms of post-transcriptional and translational regulation.

### PIWIL3 can both suppress transposons during oogenesis yet activates them during early embryogenesis

Last but not least, our study reveals that PIWIL3 plays critical and dual-functional role in TE regulation. Contrary to the canonical view of PIWI proteins as consistent TE silencers, PIWIL3 preferentially silencing a large number of TEs during oocyte maturation yet activating another large set of transposons during embryogenesis (Fig. 7b). This biphasic and opposing regulation might balance two competing needs: preserving genome integrity in the oocyte while enabling TE-derived regulatory elements essential for zygotic genome activation (ZGA). The collapse of PIWIL3-bound piRNAs—which constitute over 90% of small RNAs in oocytes—destabilizes TE networks, resulting in transcriptional disarray and developmental arrest.

In conclusion, our findings reveal that PIWIL3 is essential for piRNA biogenesis (Fig. 8, pathway 1), for the post-transcriptional regulation of gene (Fig. 8, pathway 2), and a dual role in stage-specific repression and activation of TE expression (Fig. 8, pathway 3). These molecular functions play critical roles in oogenesis and early embryogenesis. These findings redefine PIWIL3 as a central orchestrator of female fertility. They represent a significant conceptual advance beyond traditional rodent-centric paradigms. By uncovering the comprehensive roles of PIWIL3 in regulating transcriptomic, proteomic, and piRNA-mediated regulatory networks, as well as its essential function in oogenesis and embryogenesis, this work positions rabbits as a critical translational model for human reproductive biology, paving the way for innovative diagnostic and therapeutic interventions in infertility.

**Fig. 8.**
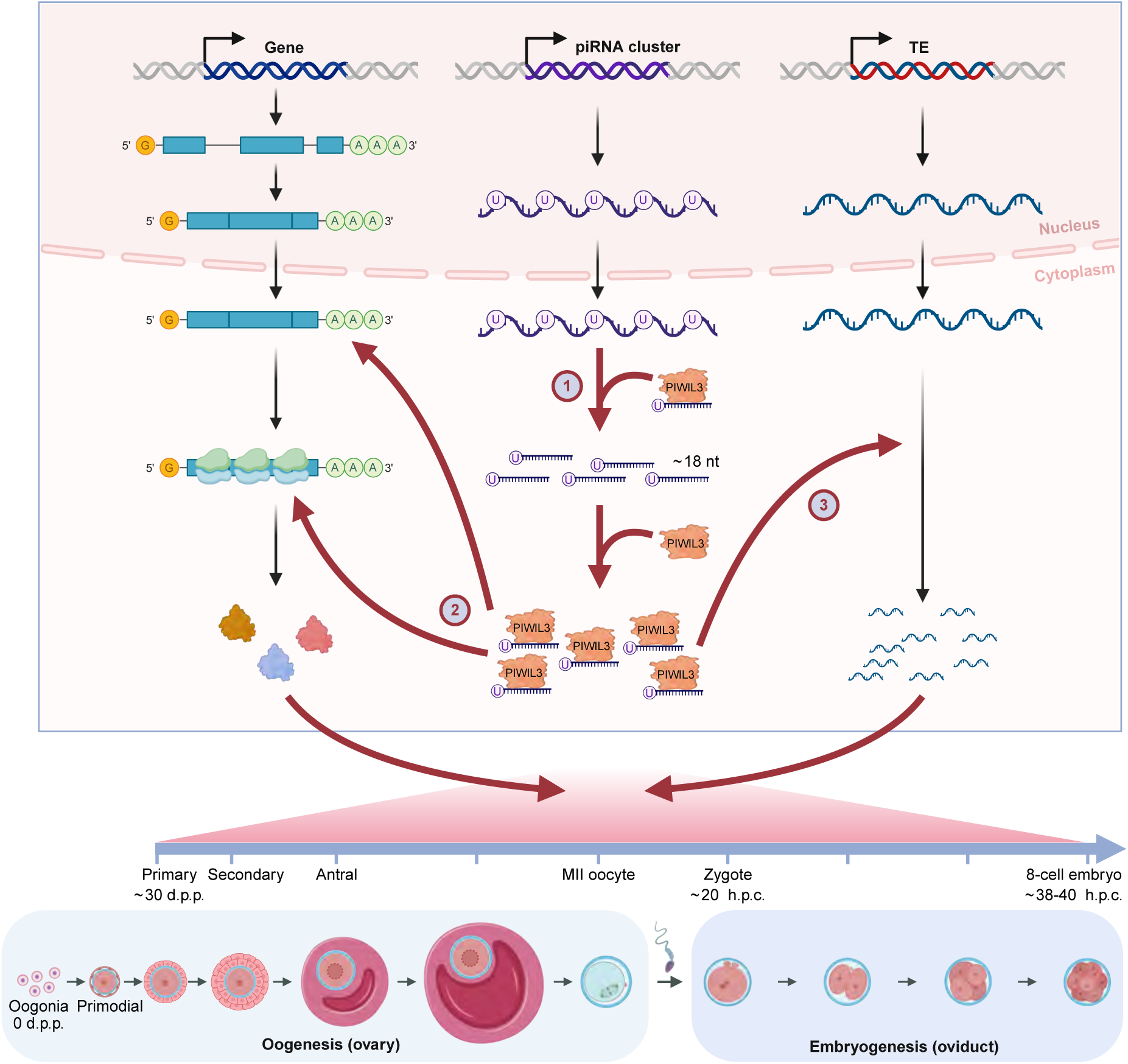
A model of PIWIL3 function in rabbit oogenesis and embryogenesis. This model illustrates the logical connections between the diverse roles of PIWIL3 in 18-nt piRNA biogenesis, in regulating a broad spectrum of genes at both mRNA and protein levels, and in its dual-function involvement in TE regulation-processes collectively essential for normal rabbit oogenesis and as a maternal effect protein for embryogenesis.

## MATERIALS AND METHODS

### Animals and ethics statement

New Zealand white rabbits were obtained from the Laboratory Animal Center of Jilin University (Changchun, China) and Shanghai Jiao Tong University (Shanghai, China). All animal studies were conducted according to the guidelines for the Care and Use of Experimental Animals approved by the Institutional Animal Care and Use Committee (IACUC) of ShanghaiTech University.

### Generation of PIWIL3-deficient rabbit

One single sgRNA for *PIWIL3*^1^*^/^*^1^ strain and one pair of sgRNAs for *PIWIL3*^2^*^/^*^2^ strain were designed to knock out PIWIL3 according to a previous protocol ^53^. Briefly, the gRNA sequences were cloned into pUC57-Simple-gRNA (Addgene ID 51306) vector and transcribed *in vitro*. To produce SpCas9 mRNA, the PCS2+Cas9 (Addgene ID 122,948) plasmid was linearized with NotI restriction digestion and used as a template for *in vitro* transcription. Microinjection of pronuclear-stage embryos was performed according to our established method ^54^. Briefly, a mixture of Cas9 (200 ng/µl) and sgRNA (50 ng/µl) mRNA was co-injected into the cytoplasm of pronuclear-stage rabbit zygotes and then transferred into the recipient rabbits, Recipients were allowed to naturally deliver at around day 30 and raise their pups.

### Rabbit genotyping

Genomic DNA was isolated from ear clips of newborn rabbits using the QuickExtract DNA Extraction Solution (Lucigen, QE09050) for PCR genotyping, followed by Sanger sequencing. All primers used for detection are detailed in Supplementary Table 1.

### Collection of GV, GVBD, and MII oocytes, zygotes, as well as 2-cell, 4-cell, 8-cell and 16-cell embryos

New Zealand white rabbits GV and GVBD oocytes were collected from ovaries as described previously ^55^. MII oocytes, zygotes, 2-cell, 4-cell, 8-cell and 16-cell embryos were collected through surgical oviduct flushing from donors after superovulation treatment, also as described previously ^56^. Briefly, female New Zealand white rabbits (6-8 months old) in proestrus were administered six intramuscular injections of 50 IU FSH (NSHF, Follicle Stimulating Hormone for Injection) at 12-hour intervals, followed by a single intravenous injection of 100 IU HCG (NSHF, Chorionic Gonadotrophin for Injection) via the marginal ear vein. Ovulation was induced after HCG administration, either through vaginal stimulation for MII oocyte collection or natural mating for embryo collection. The rabbits were euthanized 18 hours post-induction, and oviducts were flushed with 10 mL of Embryo manipulation medium (Medium 199, Hank’s supplemented with 10% fetal bovine serum) to collect MII oocytes or pronuclear-stage zygotes. The zygotes were transferred into embryo culture medium (containing EBSS, supplemented with 1% nonessential amino acids and 2% essential amino acids. 0.0146% W/V glutamine, 0.0044% W/V sodium pyruvate and 10% of fetal bovine serum) and cultured at 38.5 °C under 5% CO2. Embryos at the 2-cell, 4-cell, 8-cell, and 16-cell stages were collected at 26, 36, 46, and 70 hours post-coitum, respectively.

### Anti-PIWIL3 antibody production

Monoclonal anti-PIWIL3 antibodies were produced in mouse by Shanghai Youke Biotechnology Co., Ltd (Shanghai, China). The peptide NINWDQYPEDTFKRADGSVVTFVDYYMERYRECITDLNQPLLSSQGKWKKGQQDIPREPILLVPELCYLTGLTDPMRKNSRIMRDLALHTRLGPQRRQNELKEFIKAVQQDQTVQDELQLWDLKFDSNFLTFSGRVMDEVKVHQPRGWFEAIKNQAEWMRGTKTRMLSVETVNQWTLMYSTRCSEAADKLLDILSKIAPDMGMTISKPDKEIITDEFETYISRLKKLDLNQKQMVVCVLPNDKKDRYDD was chosen as the antigen.

### RNA extraction and real time PCR

Total RNA from various tissues and oocytes was isolated using TRIzol (Invitrogen, Cat#: 15596026CN) and cDNAs were prepared from 500 ng of total RNA with the HiScript III 1^st^ Strand cDNA Synthesis Kit (+ gDNA wiper, Vazyme, Cat#: R312-02). qRT-PCR reactions were performed according to the protocol of the iTaq Univer SYBR Green Supermix (Bio-Rad, Cat#: 1725125). The sequence of primers used for mRNA are list in Supplementary information, Table S1.

### 5’ and 3’ RACE

Total RNA for RACE experiment was extracted from the ovaries of 2-month-old rabbits using TRIzol (Invitrogen, Cat#: 15596026CN), 5’ and 3’ RACE was performed using the HiScript-TS 5’/3’ RACE Kit (Vazyme, Cat#: RA101) according to the protocol of manufacturer. The PCR-amplified fragments were cloned into the pCE2 TA/Blunt-Zero vector using the 5 min TA/Blunt-Zero Cloning Kit (Vazyme, Cat#: C601). Gene-specific primers were designed with RACE Primer designer (Vazyme) and listed in Supplementary information, Table S1.

### Protein extraction and Western Blot

For Figure 1J, total proteins from 20 MII oocytes were extracted by 1X NEBuffer for PMP (50 mM HEPES, 100 mM NaCl, 2 mM DTT, 0.01% Brij-35, pH 7.5, supplemented with 1mM MnCl2) supplemented with either Phosphatase Inhibitor (Roche, Cat#: 4906837001) or Lambda Protein Phosphatase (NEB, Cat#: P0753S) according to the protocol of the manufacturer. For Extended Data Figure 2D, total proteins from 20 MII oocytes were extracted by RIPA buffer (BioVision, Cat#: 2114-100) supplemented with protease inhibitor (Roche, Cat#: 11873580001) and Phosphatase Inhibitor (Roche, Cat#: 4906837001) according to the protocol of the manufacturer. Western blot was performed according to our previous method ^57^. The following antibodies were used: the anti-PIWIL3 monoclonal antibody was used at 1:500 dilution and anit-GAPDH (14C10) polyclonal antibody (Cell Signaling Technology, Cat#: 2118S) was used at 1:1000 dilution.

### Immunofluorescence and Hematoxylin-eosin staining

Formalin-fixed, paraffin-embedded ovaries from rabbits were used for both immunofluorescence and histological analysis. Formalin-fixed, paraffin-embedded testes from rabbits were used for histological analysis. Immunofluorescence staining was performed using the following primary antibodies: Anti-PIWIL3 antibody (1:200), Anti-DDX4/MVH antibody (used at 1 µg/ml final concentration, Abcam, Cat#: ab13840), Anti-AIF antibody(E20)-Mitochondrial Marker (1:500, Abcam, Cat#: ab32516), Oct-3/4 Antibody H-134 (1:100, Santa Cruz Biotechnology Inc, Cat#: sc-9081). The following secondary antibodies were used: Goat anti-rabbit IgG (H+L) highly cross-adsorbed secondary antibody conjugated to Alexa Fluor™ Plus 555 (1:500, Thermo Fisher scientific, Cat#: A32732) and goat anti-mouse IgG (H+L) highly cross-adsorbed secondary antibody conjugated to Alexa Fluor™ Plus 647 (1:500, Thermo Fisher scientific, Cat#: A32728). Fluorescence microscopic images were acquired using Zeiss LSM 710 confocal microscope and analyzed using ZEN (version 2012). Hematoxylin-eosin staining was performed using a Hematoxylin and Eosin Staining Kit (Yeasen, Cat#: 60524ES60) based on the manufacturer’s instructions. Hematoxylin-eosin staining images were captured using TissueFAXS PLUS Panoramic Tissue Cell Quantitative Analysis System.

### Single-cell mRNA library construction

It was performed according to the previously described Smart-seq3 method with some modifications ^58^. In brief, single oocyte or embryo was incubated with lysis buffer mix (5% (v/v) Poly-ethylene Glycol 8000, 0.1% (v/v) Triton X-100, 0.5 U/µL RNase Inhibitor, 0.5 µM OligodT30VN, 0.5 mM dNTPs) at 72 °C for 10 minutes to release RNA. After reverse transcription and PCR pre-amplification, cDNA products were purified using the Agencourt Ampure XP beads at 0.6:1 ratio (Beckman Coulter, Cat#: A63881). Library preparation was performed using 0.5 ng of purified cDNA with the Nextera XT DNA Library Preparation Kit (illumina, Cat#: 15032785), employing Tn5 transposase for tagmentation. The amplified libraries were purified with Agencourt AMPure XP beads at a 0.6:1 (bead:sample) ratio and sequenced in 2 × 150 bp paired-end mode on an Illumina HiSeq 2500 platform.

### Processing of single-cell mRNA sequencing data

The zUMIs pipeline was used to process Smart-seq3 data ^59^. Reads mapped to genes or transposons were counted by supplying either gene annotations or transposon annotations in input yaml files. Down-sampled and normalized count tables were reported in dgecounts.rds for each feature type (exons, introns, intron+exon). Differentially expressed genes and transposons meeting the (FDR < 0.05, |fold change| ≥ 1.5) criteria were selected for downstream analysis. Pseudotime analysis was performed by monocle3 (http://cole-trapnell-lab.github.io/monocle3/) with the default parameters.

### LC-MS/MS analysis

Each single oocyte or embryo was added first with SDT buffer (4% SDS, 100 mM Tris-HCl, pH 7.6) and then with DTT (Bio-Rad, Cat#: 161-0404) to the final concentration of 40 mM. The samples were mixed at 600 rpm for 1.5 hours at 37 °C to lyse cells. After the samples cooled to room temperature, Iodoacetamide (Bio-Rad, Cat#: 163-2109) was added into the mixture to the final concentration of 20 mM to block reduced cysteine residues and the samples were incubated for 30 minutes in darkness. Next, the samples were transferred to the filters (Microcon units, 10 kDa), washed with 100 µl UA buffer (8M urea, 150 mM Tris-HCl, pH 8.0) three times and then with 100 µl 25 mM NH4HCO3 buffer twice. Finally, trypsin was added to the samples (the trypsin: protein (wt/wt) ratio was 1:50) and incubated at 37 °C for 15-18 hours (overnight). The resulting peptides were collected as filtrates, desalted on C18 Cartridges (Empore™ SPE Cartridges MCX, 30UM, waters), concentrated by vacuum centrifugation, and reconstituted in 40 µl of 0.1% (v/v) formic acid. The peptide content was estimated by UV light spectral absorbance at 280 nm. Indexed retention time calibration peptides were spiked into the sample. The peptides from each sample were analyzed by Orbitrap^TM^ Astral^TM^ mass spectrometer (Thermo Scientific) connected to an Vanquish Neo system liquid chromatography (Thermo Scientific) in the data-independent acquisition (DIA) mode. Precursor ions were scanned at a mass range of 380-980 m/z, MS1 resolution was 240000 at 200 m/z, Normalized AGC Target: 500%, Maximum IT: 5 ms. 299 windows were set for DIA mode in MS2 scanning, Isolation Window: 2 m/z, HCD Collision Energy: 25 ev, Normalized AGC Target: 500%, Maximum IT: 3 ms.

### Processing of LC–MS/MS data

DIA data was analyzed with DIA-NN 1.8.1. Main software parameters were set as follows: enzyme is trypsin, max missed cleavage is 1, fixed modification is carbamidomethyl(C), dynamic modification is oxidation(M) and acetyl (Protein N-term). All reported data were based on 99% confidence for protein identification as determined by false discovery rate (FDR) ≤ 1%.

### Single-cell small RNA library construction

Small RNA libraries from single oocyte or embryo were prepared following established protocols ^60^ with several optimizations. Briefly, individual specimens underwent thermal treatment at 72°C for 3 minutes to enhance RNA liberation and structural relaxation. Post 3’-adapter ligation, an optimized enzymatic digestion system (5 U λ exonuclease combined with 25 U 5’ deadenylase) demonstrated superior efficiency compared to conventional RecJf/deadenylase mixtures in eliminating excess adapters. Subsequent 5’-adapter ligation preceded reverse transcription to generate cDNA templates. A two-phase PCR amplification strategy was implemented, utilizing 1 µl of primary amplification product for final library enrichment. Quality-controlled libraries were size-selected through 6% denaturing polyacrylamide gel electrophoresis, specifically targeting 140-200 bp fragments. Final paired-end sequencing (2×150 bp) was conducted on the BGI-T7 platform (BGI-Shenzhen).

### Small-RNA spike-in information

To normalize between different data sets, six synthetic small-RNA spike-in controls (three unmethylated and three methylated variants) were commercially synthesized (Integrated DNA Technologies) and combined into a mixture as described in previous^60^. A defined quantity (5 × 10^-6^ pmol) of this spike-in cocktail was introduced into each reaction system during single oocyte/embryo processing. Detailed nucleotide sequences of these synthetic RNA standards were obtained from a previous study ^60^.

### Processing of single-cell small RNA sequencing data

Small RNA sequences were processed with TrimGalore (version 0.6.10; http://www.bioinformatics.babraham.ac.uk/projects/trim_galore/) with the default adapter trimming mode to auto-detect adapters in the 150bp long sequencing reads. A size cutoff using 15 to 42 nt was applied to retain small RNA reads of suitable length. Subsequently trimmed reads were mapped onto the rabbit genome using Bowtie1 (version 1.2.2) with the parameters-v 0 -n 0 -l 18 -k 1, allowing perfect matches and reporting 1 alignment. Genome-mapped reads were sequentially mapped to the libraries of miRNA, tRNA, rRNA, snRNA, snoRNA, piRNA, and repetitive elements. The reads in each library were normalized to the counts of exogenous spike-ins. The raw read counts for both the spike-ins and the total reads are provided in Supplementary information, Table S2.

### Mapping of PIWIL3-bound piRNAs to mRNAs and TEs

PIWIL3-bound piRNAs were mapped to the reverse complementary sequence of differentially expressed mRNAs, proteins, or TEs using Bowtie1 (v1.2.2) with parameters -n 0 -l 15 -a (perfect matches) or -n 1 -l 15 -a (allowing one mismatch).

### Weighted gene co-expression network analysis (WGCNA)

WGCNA was performed on single-cell mRNA, transposon and protein expression data using the WGCNA R package (v1.72; https://cran.r-project.org/web/packages/WGCNA/index.html). The analysis was conducted with the following parameters: Network type: Signed, Soft-thresholding power (β): 9 for mRNA and transposon data, 8 for protein data (determined by scale-free topology criterion with R² > 0.85), Minimum module size: 30 genes, Module merging threshold: 0.25, Dynamic tree cutting with deepSplit = 2, Intramodular connectivity (kME) > 0.8, All reported modules met significance criteria: module-trait correlation |r| > 0.5, P < 0.01.

### Source of genome annotations

The annotation of the New Zealand white rabbit genome was downloaded from UCSC genome annotation database (https://genome.ucsc.edu/index.html) for the Apr. 2009 (Broad/oryCun2) assembly of the rabbit genome (oryCun2, Broad Institute oryCun2 (NCBI project 12819, AAGW00000000)). Gene annotations were also downloaded from UCSC. Functional RNA from New Zealand white rabbit was retrieved from the following databases: miRNAs were from genome annotation (Ensembl GTF); tRNAs were from GtRNAdb (https://gtrnadb.ucsc.edu/); snoRNAs and snRNAs were from RNAcentral v17 (https://rnacentral.org); rRNAs were compiled from both genome annotation (Ensembl GTF) and RNAcentral. Transposon annotations were downloaded from RepeatMasker (https://www.repeatmasker.org/). piRNA clusters were annotated and classified using proTRAC (v2.4.2) based on genome-aligned small RNA reads (15-42 nt) retained after miRNA depletion in MII-stage rabbit oocytes^61^. Commands for proTRAC were: “perl proTRAC_2.4.3.pl -map ${map_files} -genome oryCun2.fa -geneset oryCun2.ensGene.sorted.gtf -repeatmasker oryCun2.RepeatMasker_original -pimin 15 -pimax 34”. $mapfile was the mapped output using sRNAmapper (version 1.0.4) ^62^. Repeat-derived small RNA libraries were obtained from the RepeatMasker annotation in UCSC genome browser. Any overlapping clusters were merged into one cluster. piRNA candidates were identified by mapping onto rabbit piRNA clusters.

### Genomic annotation of piRNA

piRNA sequences were first mapped to previously defined piRNA clusters using Bowtie with the parameters -v 0 -n 0 -l 18 -a, allowing perfect matches and reporting all possible alignments. The resulting BAM files were converted to BED format, with coordinates adjusted to genomic positions. For each piRNA, a fractional weight of 1/n was assigned to each mapped location, where n is the total number of mapping sites for that sequence.

The weighted BED file was then intersected with various genomic annotation datasets, including RepeatMasker elements, protein-coding genes, lncRNAs, and pseudogenes. If a locus overlapped multiple annotations, it was assigned a feature label based on the following priority: (1) repeat_region_antisense, (2) repeat_region_sense, (3)

protein_coding_sense, (4) lncRNA_sense, (5) pseudogenes_sense, and (6) processed_pseudogenes_sense. Additionally, we annotated piRNAs based on specific transposon subfamilies and sub-genic regions within protein-coding genes (i.e., CDS, 3′UTR, and 5′UTR), using the same weighted BED approach. For each annotation category, the total count was computed by summing the assigned weights across all mapped piRNAs.

### RNA immunoprecipitation (RIP)

For each condition, 200 GV/GVBD rabbit oocytes were lysed in RIP lysis buffer according to our established method ^63^. Dynabeads Protein G (Thermo Fisher Scientific, Cat#: 10004D) were washed once with lysis buffer and coupled with 5 µg of homemade anti-PIWIL3 or mouse IgG (Millipore, Cat#: 12-371) antibodies at 4 °C for 4 hours. Oocyte lysates were mixed with antibody-coupled beads and rotated gently at 4 °C overnight. The beads were then washed twice with RIP wash buffer and two times with RIP high wash buffer as described in our previously method ^64^. The RNA-containing supernatant was recovered using a magnetic holder and processed for both single-cell small RNA and smart-seq3 library construction, as described above.

### Statistics and reproducibility

For comparisons between data groups, two-tailed Student’s T-tests were used. Exact P values are shown in the figures. Data were presented as the mean ± SD or mean ± SEM and P < 0.05 was considered significant. Statistical analysis was performed using GraphPad Prism 10.1.2 software. Pearson’s correlation analysis was used to calculate the regression and correlation between two groups. As indicated in the figure legends, all assays were performed in three biological replicates unless stated otherwise. Representative micrographs and Western blot shown in figures were repeated three times independently with similar results.

## Data availability

The accession number for the mRNA and small RNA data reported in this paper is GEO (https://www.ncbi.nlm.nih.gov/geo/): PRJNA1241353. The mass spectrometry proteomics data have been deposited to the ProteomeXchange Consortium (https://proteomecentral.proteomexchange.org) with the dataset identifier PXD063649. Source data are provided with this paper.

## ACKNOWLEDGEMENTS

We thank Prof. Ligang Wu for providing the Single-cell CAS-seq system. We acknowledge Ms. Zhangyue Song of the Biomedical Big Data Platform at the Shanghai Institute for Advanced Immunochemical Studies, ShanghaiTech University, for her support. We are also grateful to the Animal Core Facility and the Molecular Imaging Core Facility at the School of Life Science and Technology, ShanghaiTech University. We thank the ShanghaiTech High Performance Computing Platform for providing computational resources. Additionally, we sincerely appreciate the valuable comments and suggestions from Prof. Tian Chi, Prof. Enzhi Shen, Prof. Liye Zhang, and Dr. Chen Wang. We acknowledge BioRender (https://www.biorender.com/) for providing the tools used to create figures in this manuscript, under Academic License HU28BQ3T11.

This work was supported by start-up funding from ShanghaiTech University and Shanghai Natural Science Foundation (25ZR1401254 to S.S.).

## AUTHOR CONTRIBUTIONS

H.L. conceived the project. H.L. (1.2015-5.2022) and S.S. (6.2022-2025) supervised the study, Y.G., S.S., L.L. performed experiments. S.G. created *PIWIL3*^1^ mutant, Y.Q. and Z.L. created *PIWIL3*^2^ mutant. Y.G., S.S., T.L., and H.L. analyzed data. S.S., Y.G., and H.L. wrote the manuscript.

## CONFLICT OF INTEREST

The authors declare no competing interests.

---

**Fig. S1.**
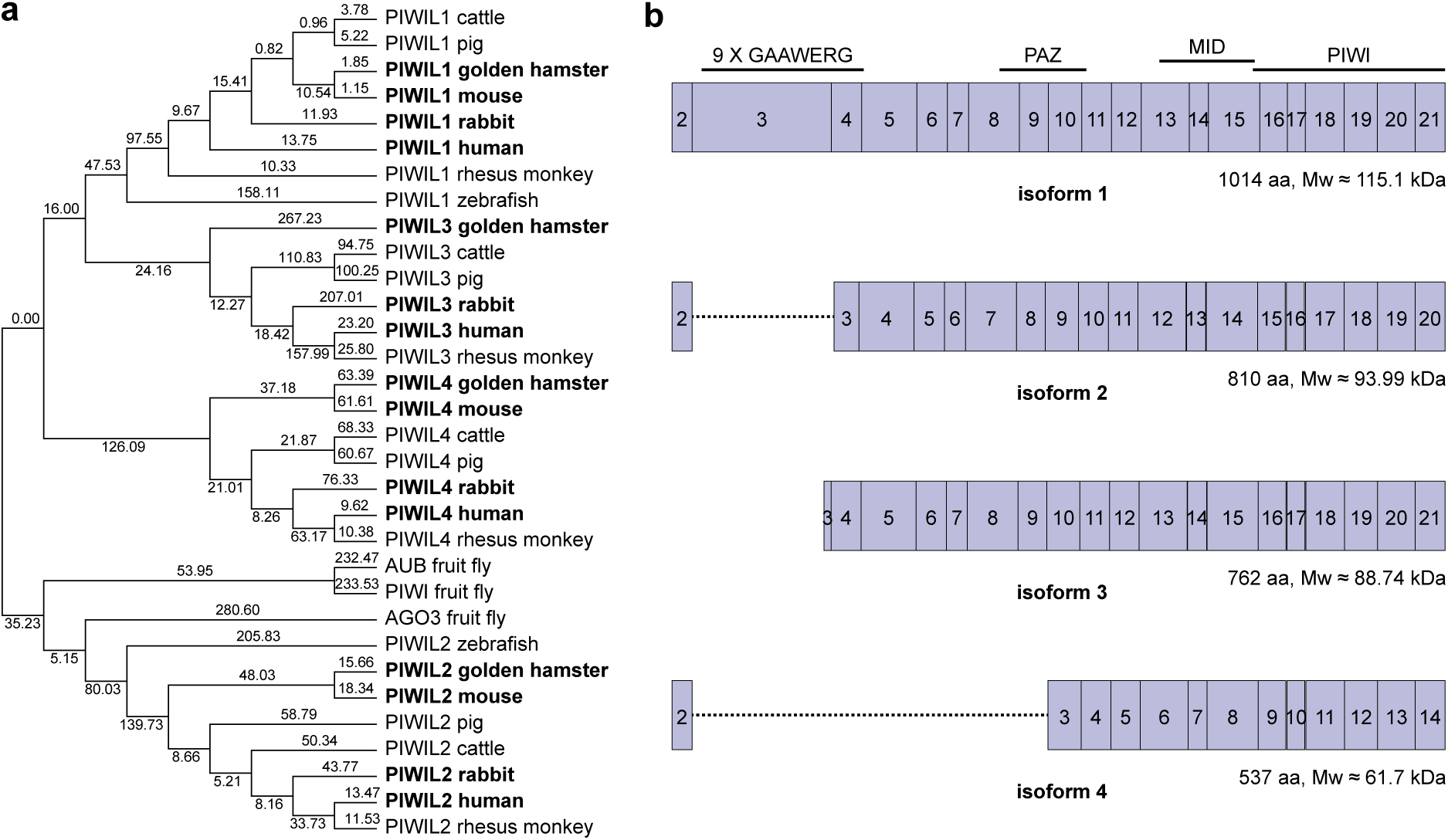
Characterization of PIWIL3 transcripts in rabbit. **a** Phylogenetic tree illustrating the relationships of *PIWI* genes across nine species. The numbers on the tree represent the estimated divergence times in millions of years. **b** Nucleotide sequence alignment of four *PIWIL3* transcript isoforms identified in rabbits.

**Fig. S2.**
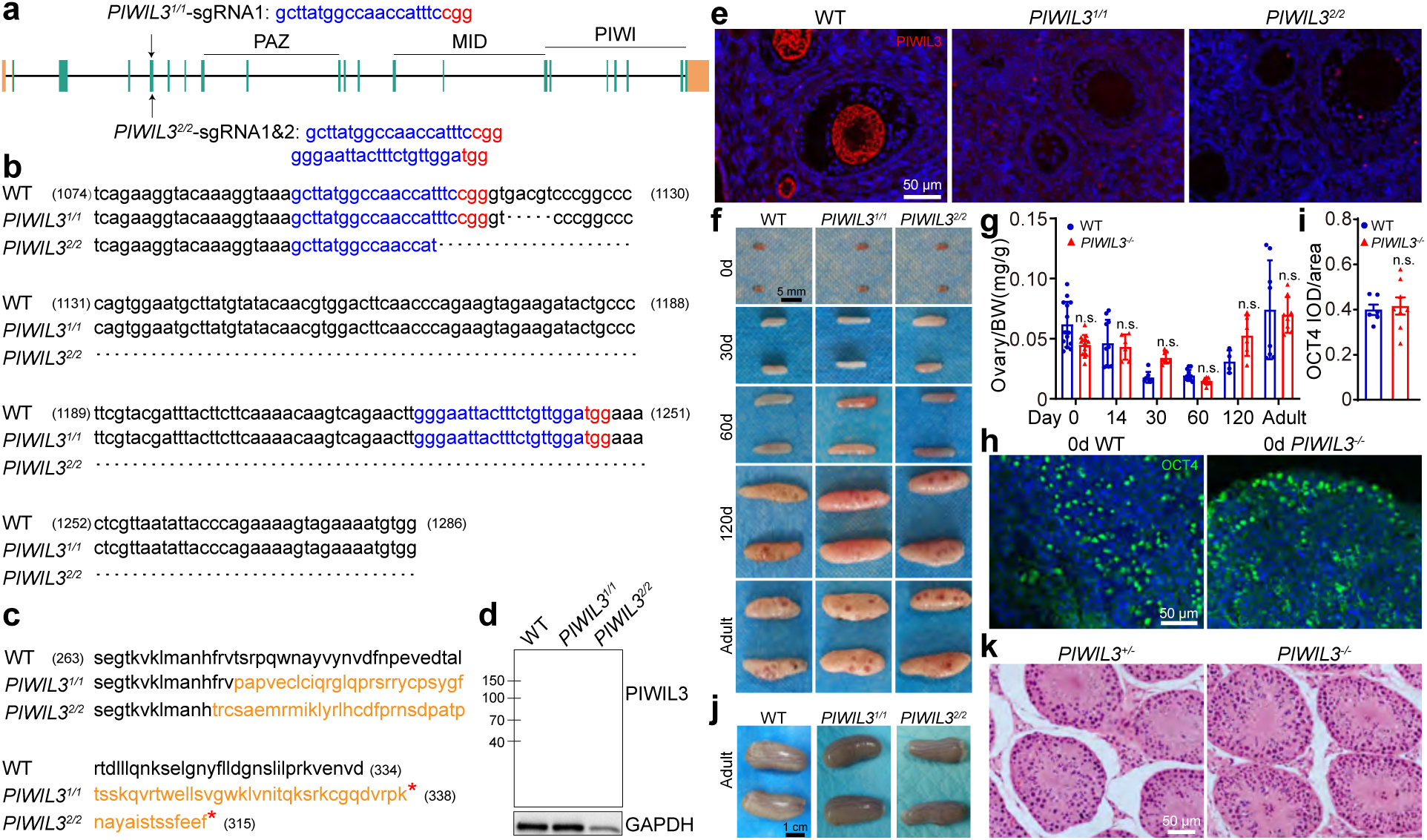
Generation of *PIWIL3* mutant rabbits. **a** Structure of the rabbit *PIWIL3* gene and sgRNA target sites. Orange regions represent untranslated regions (UTRs), while green regions denote protein-coding sequences. Two mutant strains were generated using CRISPR-Cas9: Mutant 1 (*PIWIL3*^1^*^/^*^1^) was targeted with sgRNA1 on Exon 5, and Mutant 2 (*PIWIL3*^2^*^/^*^2^) was targeted with sgRNA1 and sgRNA2 on Exon 5. **b** Nucleotide sequences of the *PIWIL3* target site in WT and mutant strains (*PIWIL3*^1^*^/^*^1^ and *PIWIL3*^2^*^/^*^2^). **c** Amino acid sequences of the PIWIL3 target site in WT and mutant strains (*PIWIL3*^1^*^/^*^1^ and *PIWIL3*^2^*^/^*^2^). Sequences highlighted in orange indicate altered sequences, and red stars mark premature stop codons. **d** Western blots showing the loss of PIWIL3 expression in *PIWIL3*-deficient rabbit oocytes. GAPDH was used as the loading control. **e** Immunostaining analysis shows the loss of PIWIL3 expression in *PIWIL3*-deficient ovaries. Scale bars = 50 µm. The experiments presented in panels D and E were independently repeated three times with consistent results. **f** Ovaries from WT and *PIWIL3*-deficient (*PIWIL3*^1^*^/^*^1^ and *PIWIL3*^2^*^/^*^2^) rabbits at various ages (0 days, 30 days, 60 days, 120 days, and adult (7 months-2 years)). Scale bars = 5 mm. **g** Bar graph showing the quantitative analysis of ovarian weight in WT and *PIWIL3*-deficient rabbit ovaries at various ages (0 days, 14 days, 30 days, 60 days, 120 days, and adult (7 months-2 years)). **h** OCT4 immunostaining (green) of ovarian sections from 0-day WT and *PIWIL3*-deficient rabbits. OCT4 positive cells represent oogonia. Scale bars = 50 µm. **i** Bar graph showing the quantitative analysis of WT (n=6) and *PIWIL3*-deficient (n=8) OCT4 integrated optical density per unit area (IOD/area). **j** Testes from adult WT and *PIWIL3*-deficient (*PIWIL3*^1^*^/^*^1^ and *PIWIL3*^2^*^/^*^2^) rabbits. Scale bars = 1 cm. **k** H&E stained testis sections of adult control (*PIWIL3^+/-^*) and *PIWIL3*-deficient (*PIWIL3^-/-^*) rabbits. Scale bars = 50 µm.

**Fig. S3.**
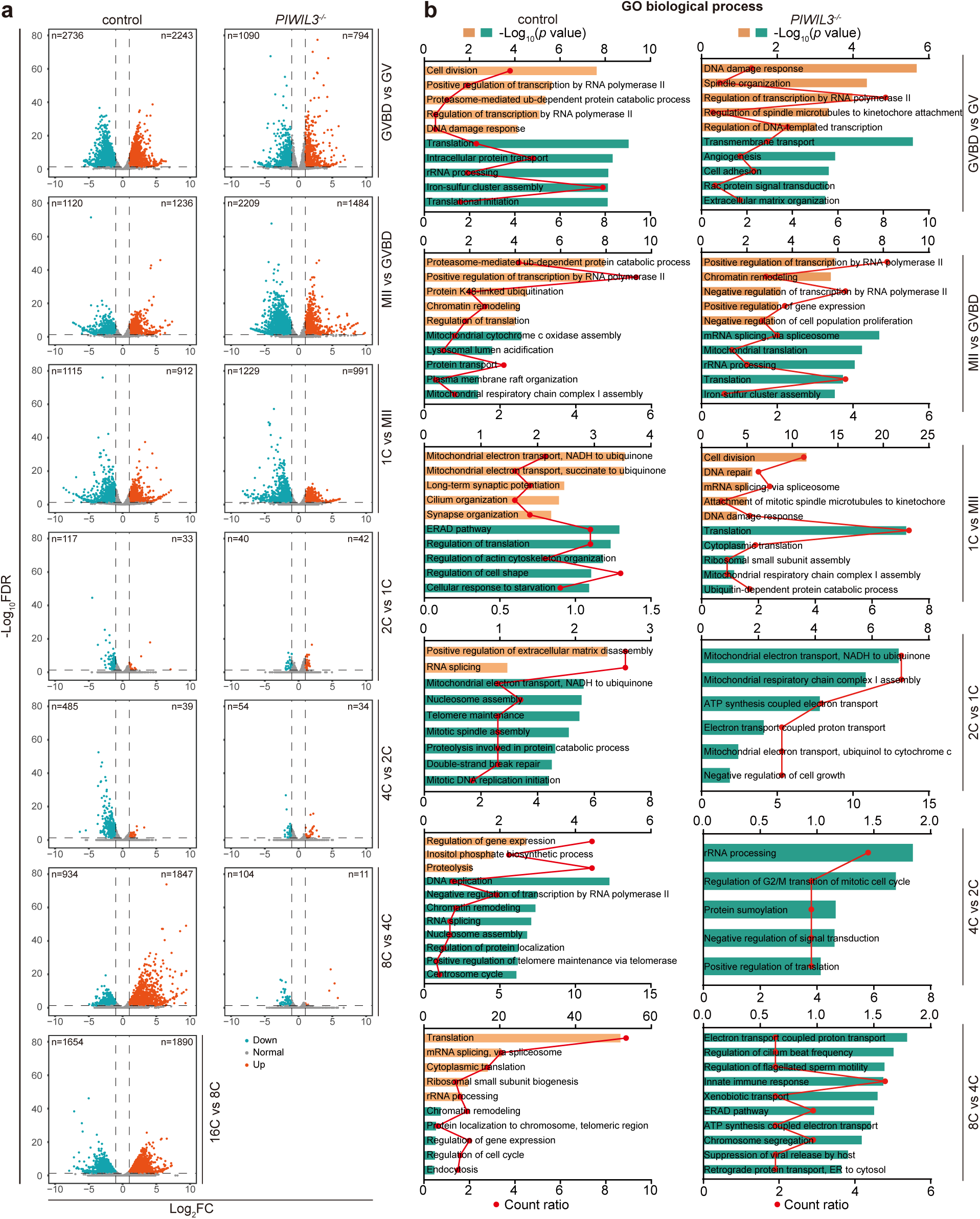
Transcriptomic profiling in *PIWIL3*-deficient oocytes and maternally *PIWIL3*-deficient embryos during the OET. **a** Volcano plots illustrating differentially expressed RNAs across the following stage comparisons in both control and *PIWIL3*-deficient samples: GVBD versus GV; MII versus GVBD; 1C versus MII; 2C versus 1C; 4C versus 2C; and 8C versus 4C. Significantly upregulated and downregulated RNAs (≥ 2-fold change; FDR ≤ 0.05) are highlighted in orange and green, respectively. **b** GO enrichment analysis of the significantly upregulated (orange) and downregulated (green) RNAs identified in panel **a** across GV-8C stage transitions. The upper x-axis shows -log₁₀(*p* value) (bar length), and the lower x-axis shows the count ratio (dots and line).

**Fig. S4.**
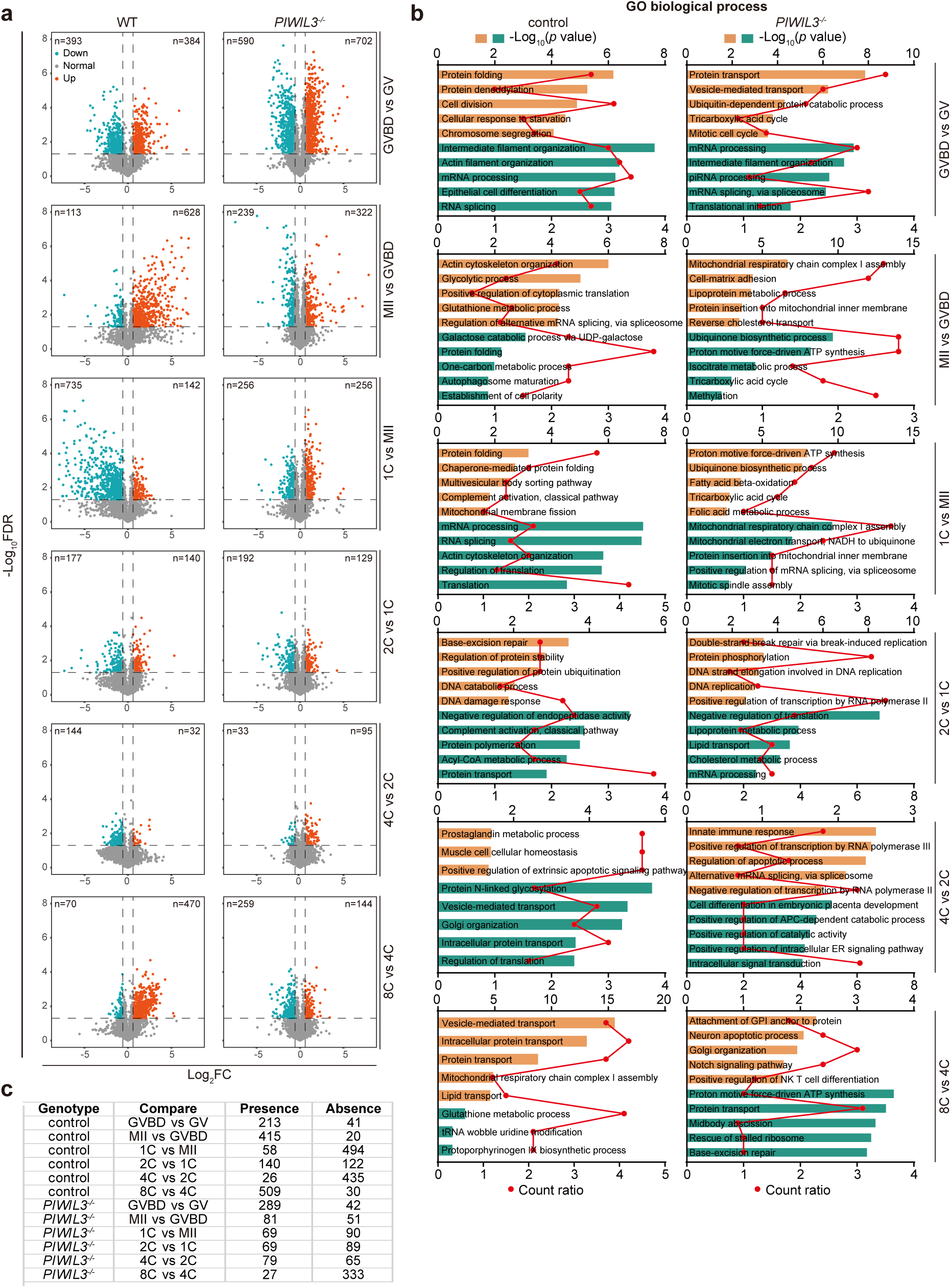
Proteomic profiling in *PIWIL3*-deficient oocytes and maternal PIWIL3-deficient embryos during OET. **a** Volcano plots illustrating differentially expressed proteins across the following stage comparisons in both control and *PIWIL3*-deficient samples: GVBD versus GV; MII versus GVBD; 1C versus MII; 2C versus 1C; 4C versus 2C; and 8C versus 4C. Significantly upregulated and downregulated RNAs (≥ 1.5-fold change; FDR ≤ 0.05) are highlighted in orange and green, respectively. **b** GO enrichment analysis of the significantly upregulated (orange) and downregulated (green) proteins identified in panel **a** across GV-8C stage transitions. The upper x-axis shows -log₁₀(*p* value) (bar length), and the lower x-axis shows the count ratio (dots and line). **c** Tables presenting the numbers of differentially present and absent genes between developmental stages in control and *PIWIL3*-deficient samples.

**Fig. S5.**
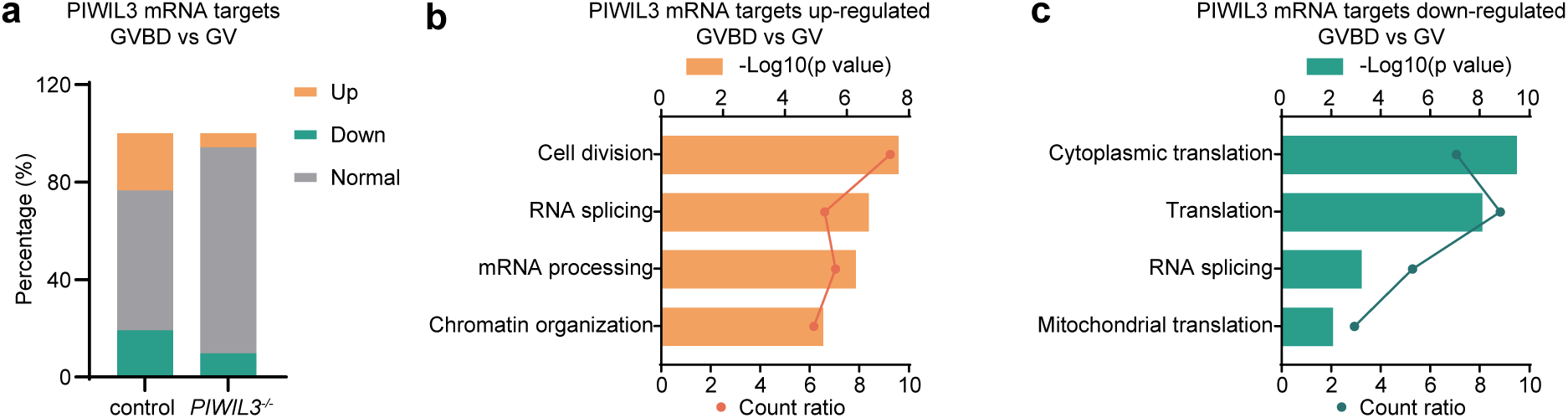
RNA dynamics of PIWIL3-bound mRNAs during the GV to GVBD oocytes transition. **a** Bar plot showing the proportions of PIWIL3-bound mRNAs that were upregulated (orange), downregulated (green), or unchanged during the GV to GVBD transition in control (left) and *PIWIL3*-deficient (*PIWIL3^-/-^*, right) oocytes. **b** GO enrichment analysis of the PIWIL3-bound mRNAs significantly upregulated in control samples (≥ 2-fold change). The upper x-axis shows -log₁₀(*p* value) (bar length), and the lower x-axis shows the count ratio (dots and line). **c** GO enrichment analysis of the PIWIL3-bound mRNAs significantly downregulated in control samples (≥ 2-fold change). Axes are as in panel **b**.

**Fig. S6.**
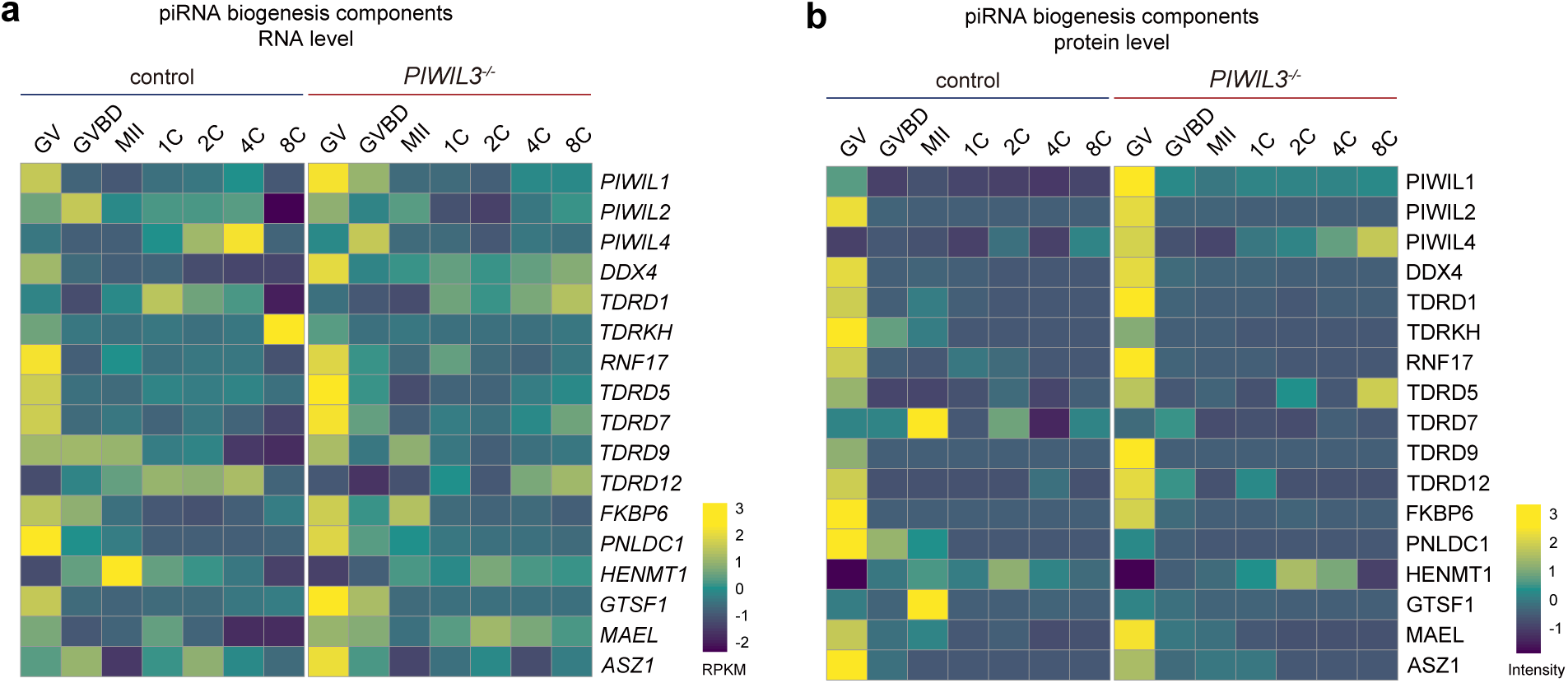
Dynamics of piRNA biogenesis components in control and *PIWIL3*-deficient oocytes and embryos. **a** Heatmap showing mRNA expression dynamics of piRNA biogenesis components from GV to 8C stages in control and *PIWIL3*-deficient (*PIWIL3^-/-^*) samples, normalized to RPKM. **b** Heatmap showing protein expression dynamics of the same genes across GV to 8C stages in control and *PIWIL3*-deficient samples, normalized to protein intensity.

**Fig. S7.**
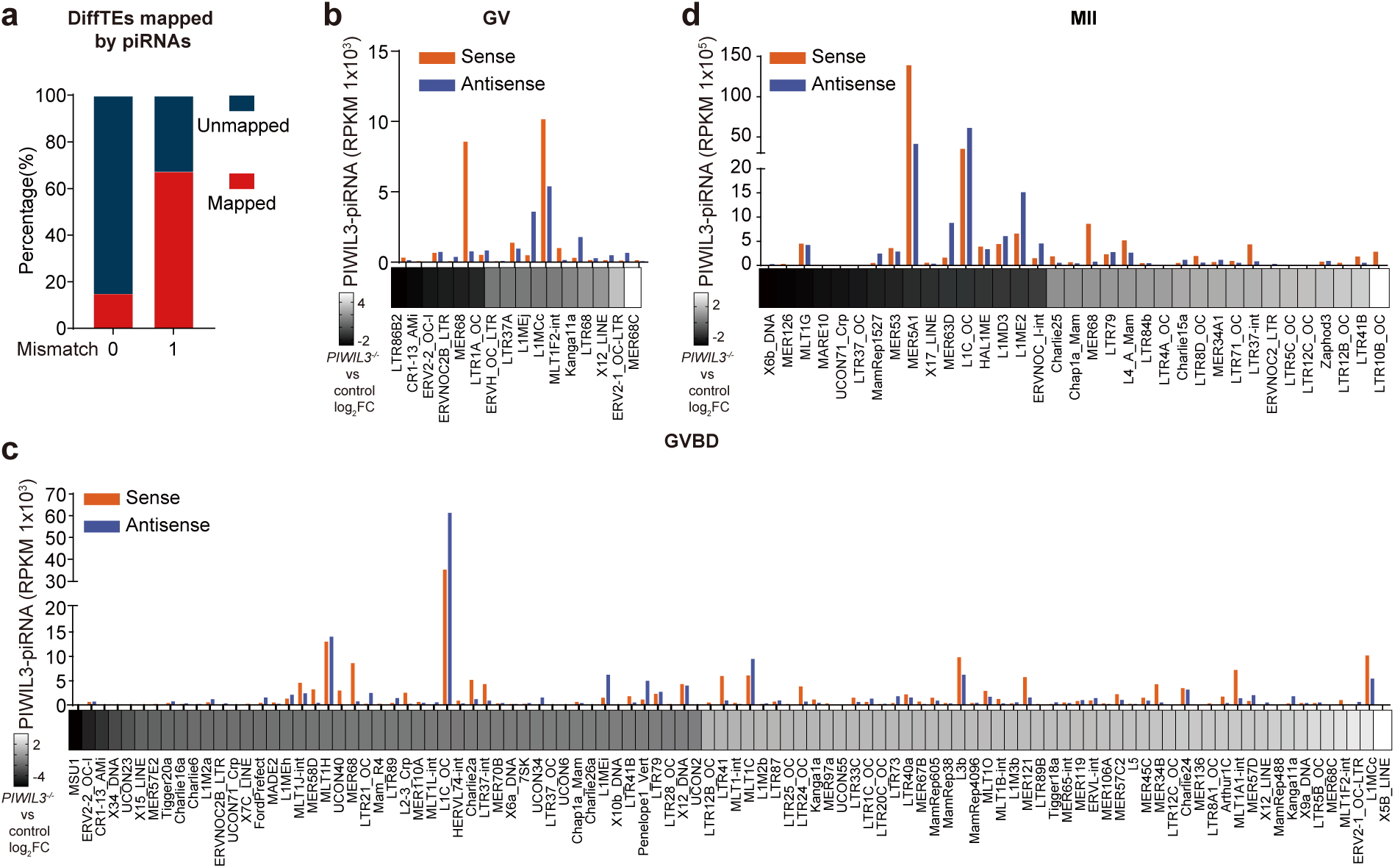
PIWIL3-piRNAs regulate TE silencing in rabbit oocytes. **a** Bar graph showing the percentage of differentially expressed TEs mapped by PIWIL3-bound piRNAs allowing 0 or 1 mismatch. **b-d** Heatmaps of log₂ fold changes in TE expression in *PIWIL3*-deficient versus control oocytes at the GV (**b**), GVBD (**c**), and MII (**d**) stages. Increased expression is shown in white and decreased expression in black. Bars above each TE indicate the abundance of sense (orange) and antisense (blue) piRNAs targeting that TE.

**Fig. S8.**
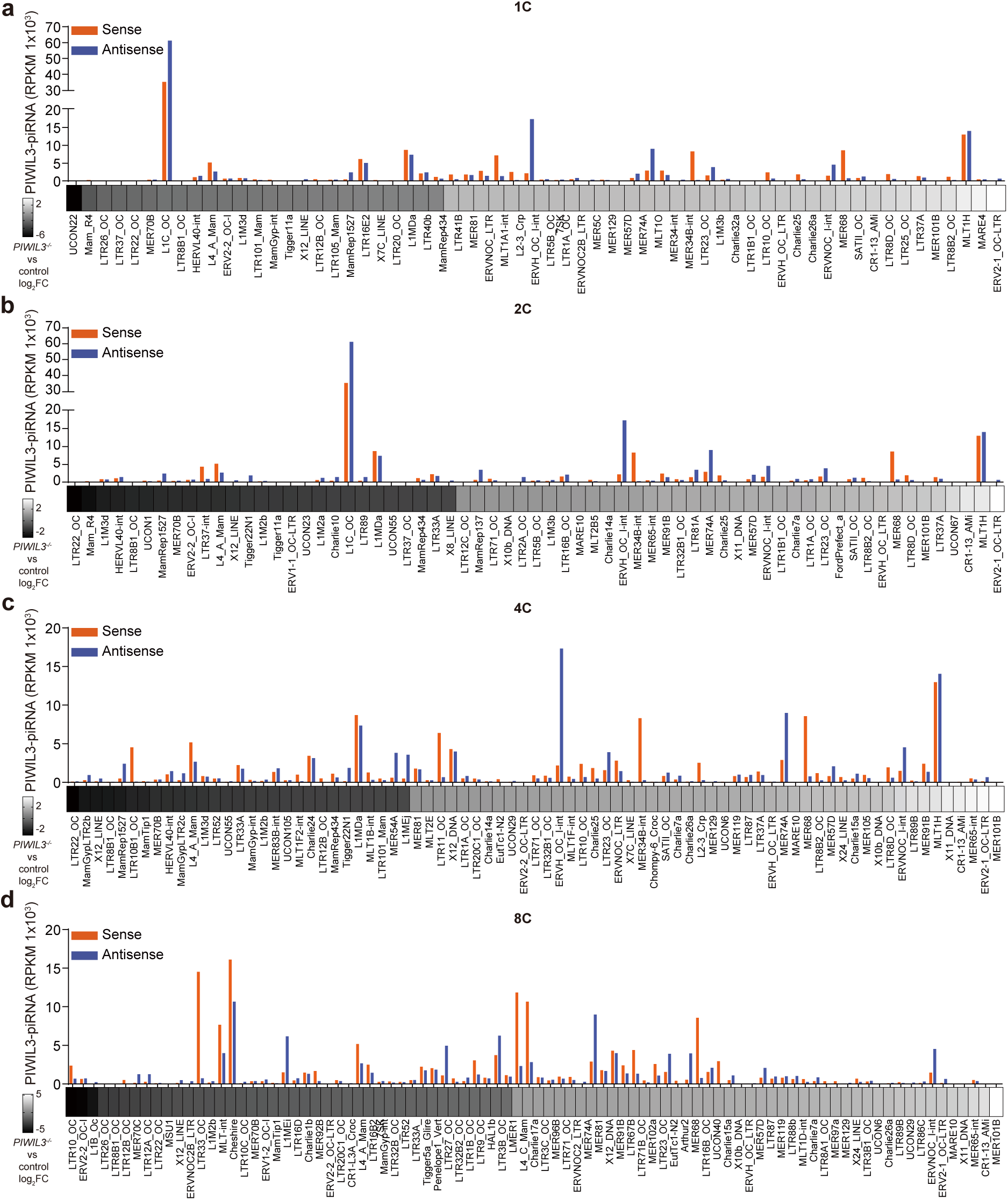
PIWIL3-piRNAs regulate TE silencing in rabbit early embryos. **a-d** Heatmaps of log₂ fold changes in TE expression in *PIWIL3*-deficient versus control embryos at the 1C (**a**), 2C (**b**), 4C (**c**), and 8C (**d**) stages. Increased expression is shown in white and decreased expression in black. Bars above each TE indicate the abundance of sense (orange) and antisense (blue) piRNAs targeting that TE.

**Fig. S9.**
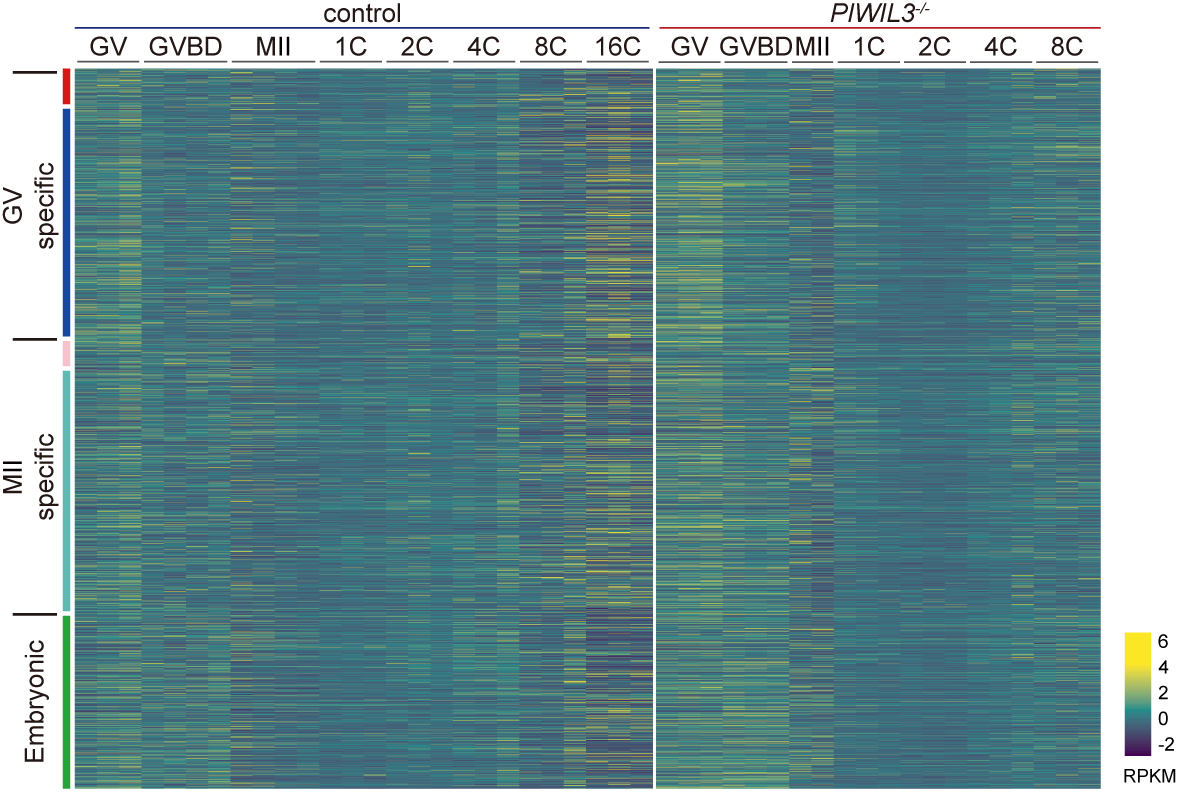
mRNA expression patterns of gene modules in control and *PIWIL3*-deficient samples. Heatmaps showing mRNA expression patterns of gene modules in control and *PIWIL3*-deficient samples across developmental stages based on protein expression patterns (see Fig. 4c), normalized to RPKM.

## Notes

### Competing Interest Statement

The authors have declared no competing interest.

